# Uncovering diversity and climatic drivers of leafhopper-parasitoid dynamics in Canada

**DOI:** 10.1101/2025.08.23.671950

**Authors:** Jordanne Jacques, Abraão Almeida Santos, Joshua Molligan, Florent Sylvestre, Nicolas Plante, Jose L. Fernandez-Triana, Joel H. Kits, Jean-François Landry, Joseph Moisan-De-Serres, Frédéric McCune, Valérie Fournier, Edel Pérez-Lopéz

**Affiliations:** Département de Phytologie, Faculté des sciences de l’agriculture et de l’alimentation, Université Laval, Québec City, Québec, G1V 0A6, Canada; Centre de Recherche et d’Innovation sur les Végétaux (CRIV), Université Laval, Québec City, Québec, G1V 0A6 Canada; Institute de Biologie Intégrative et des Systèmes (IBIS), Université Laval, Québec City, Québec, G1V 0A6, Canada; L’Institute EDS, Université Laval, Québec City, Québec, G1V 0A6, Canada; Canadian National Collection of Insects, Arachnids and Nematodes, Agriculture and Agri-Food Canada, Ottawa, ON, Canada; Laboratoire d’expertise et de diagnostic en phytoprotection, Ministère de l’Agriculture, des Pêcheries et de l’Alimentation du Québec, Québec City, Québec, G1P 3W8, Canada

**Keywords:** Biological control, *Gonatopus*, Insect migration, Genomics, Agroecosystem resilience

## Abstract

As climate change reshapes northern agroecosystems, leafhoppers (Hemiptera: Cicadellidae) are shifting their distributions, with implications for pest outbreaks and crop health. In Eastern Canada, we monitored strawberry farms from 2023 to 2024, collecting over 82,000 leafhoppers from 64 genera. Migratory species, *Empoasca fabae* and *Macrosteles quadrilineatus*, dominated captures, with sharp abundance increases above 16⍰°C and 14⍰°C, respectively, while local species declined under higher rainfall. A major finding was the first Canadian record of the corn pest *Dalbulus maidis*, a vector of multiple pathogens, likely introduced through long-distance dispersal. Insecticide applications generally failed to reduce leafhopper numbers, highlighting the limitations of current chemical control. Parasitism rates by *Gonatopus* wasps (Dryinidae), averaged ~3% but peaked in late summer at over 20%, primarily in *M. quadrilineatus*. Warmer temperatures and seasonal progression increased both parasitism probability and rates. Genomic analyses revealed at least three *Gonatopus* lineages, including the first complete mitochondrial genome for the genus from the New World, and confirmed multiple host species. We also recorded the first Canadian occurrence of *G. clavipes*. Our results demonstrate that parasitoids are active, climate-responsive, and capable of targeting dominant pest species. Together, these findings provide the first ecological and genomic baseline for leafhopper–parasitoid interactions in Canada. They point to the potential of conserving and enhancing native parasitoid populations as a foundation for climate-resilient, pesticide-free pest management strategies.

## INTRODUCTION

Ongoing climate change is reshaping insect population dynamics in North America, especially through shifts in overwintering ranges, voltinism, and seasonal phenology, with diverse outcomes depending on species and life history traits (Baker et al. 2015; Lawton et al. 2022; Plante et al. 2024; Santos et al. 2025). In high-latitude regions like Eastern Canada, warming temperatures are enabling herbivorous insects to expand their ranges northward, introducing new pressures on agroecosystems and amplifying the need for innovative pest management approaches (Plante et al. 2024; Santos et al. 2024; 2025).

Leafhoppers (Hemiptera: Cicadellidae) exemplify these emerging challenges. As both direct phloem feeders and prolific vectors of over 600 plant pathogens, including bacteria and viruses, leafhoppers are increasingly recognized as key players in climate-driven shifts affecting crop health worldwide (Cooper et al. 2023; Plante et al. 2024). Recent niche models project that eastern North America, and notably Eastern Canada, could experience substantial increases in leafhopper pest species richness under future warming scenarios (Santos et al. 2024), underscoring their value as sentinels of agroecological change (Plante et al. 2024).

While most studies have focused only on the pests, a comprehensive understanding of leafhopper dynamics demands investigation into their natural enemies, particularly parasitoids. In North America, egg parasitoids such as *Anagrus* spp. (Hymenoptera: Mymaridae) play an established role in regulating vineyard leafhopper populations (Prischmann et al. 2007; Jarrell et al. 2020; Cargnus, 2024). For instance, in British Columbia’s Okanagan Valley, at least four *Anagrus* species (*A. atomus, A. avalae, A. daanei, A. erythroneurae*) effectively parasitize leafhopper eggs during summer and winter, decreasing population size (Lowery et al. 2007). Similarly, in California, *A. daanei* has been deployed against vineyard leafhopper infestations (Jarrell et al. 2020).

By contrast, parasitoids attacking nymphal and adult leafhoppers are generally poorly studied in North America. These parasitoids include big-headed flies (Diptera: Pipunculidae) (Skevington and Marshall 1997), twisted-wing parasites (Strepsiptera: Halictophagidae) (Cook 2019), and pincer wasps of the family Dryinidae (Hymenoptera). Pincer wasps, such as specimens of the genus *Gonatopus*, remain virtually unstudied in North America despite their likely ecological role (Moya-Raygoza et al. 2006). These wasps grasp and parasitize leafhopper using their chelate protarsi, but also feed on hosts, acting as either parasitoids or predators. The wingless female of the *Gonatopus* genus is taxonomically challenging due to pronounced morphological plasticity and the near-total absence of molecular data in the Nearctic region (Tribull 2015; He et al. 2020; Virla et al. 2023). Indeed, GenBank and other sequence repositories are heavily biased toward Asian and Oceanian records, leaving North American leafhopper parasitoids largely unsequenced and complicating efforts to use DNA-based monitoring or resolve cryptic species complexes.

This knowledge gap is particularly concerning given the mounting evidence that climate change not only influences leafhopper distributions but might also increase insecticide resistance, as suggested by our recent findings (Plante et al. 2024). Conservative biological control provides a critical pathway to reduce reliance on chemical control and mitigates the environmental impacts of intensive pesticide use (Biddinger et al. 2014; Wilson et al. 2017; Jennings et al. 2017). Yet without baseline knowledge of local parasitoid communities and their climatic responses, such strategies remain speculative in northern ecosystems.

Here, we address these knowledge gaps by building on our previous work, which documented the dominance of two migratory leafhopper species, *Macrosteles quadrilineatus* (Forbes, 1885) and *Empoasca fabae* (Harris, 1841), in Eastern Canada (Plante et al. 2024). While most leafhoppers in our study area are thought to spend their entire life cycles locally, part (*M. quadrilineatus*) or all (*E. fabae*) of the populations of these species overwinter further south and perform annual migrations into our area. We further characterized the core leafhopper community composition across Eastern Canada, identifying genera consistently present across years. Additionally, we report the presence of *Dalbulus maidis* (DeLong and Wolcott), a new pest leafhopper species in Canada that, over the past decade, has been expanding its distribution while transmitting multiple plant pathogens, thereby threatening the corn industry across the Americas.

To the best of our knowledge, this is the first study of its kind in Canada, combining a molecular survey of leafhopper parasitoids with assessments of parasitism rates, the host species affected, and the taxonomy of adult parasitoids, all supported by genomic evidence. Interestingly, we uncovered an unexpected diversity of *Gonatopus* species and identified key climatic drivers shaping parasitism patterns. Our integrative approach provides foundational genomic data on North American parasitoid populations and new insights into tri-trophic interactions under climate change, paving the way for biological control strategies that can reduce pesticide dependence and enhance the resilience of northern agroecosystems.

## MATERIALS AND METHODS

### Experimental setting

Consistent with our previous survey, this study was carried out in strawberry farms across five regions of Québec, Canada: Capitale-Nationale, Chaudière-Appalaches, Mauricie, Centre-du-Québec, and Montérégie (**Fig. 1a, Table S1**). Whenever possible, two fields (sites) per farm were surveyed, depending on site availability. In 2023, leafhoppers were collected from 11 farms across these regions, totaling 18 sites. In 2024, sampling took place in 10 farms, covering 16 sites, as the Centre-du-Québec region was not included that year.

**Figure 1.**
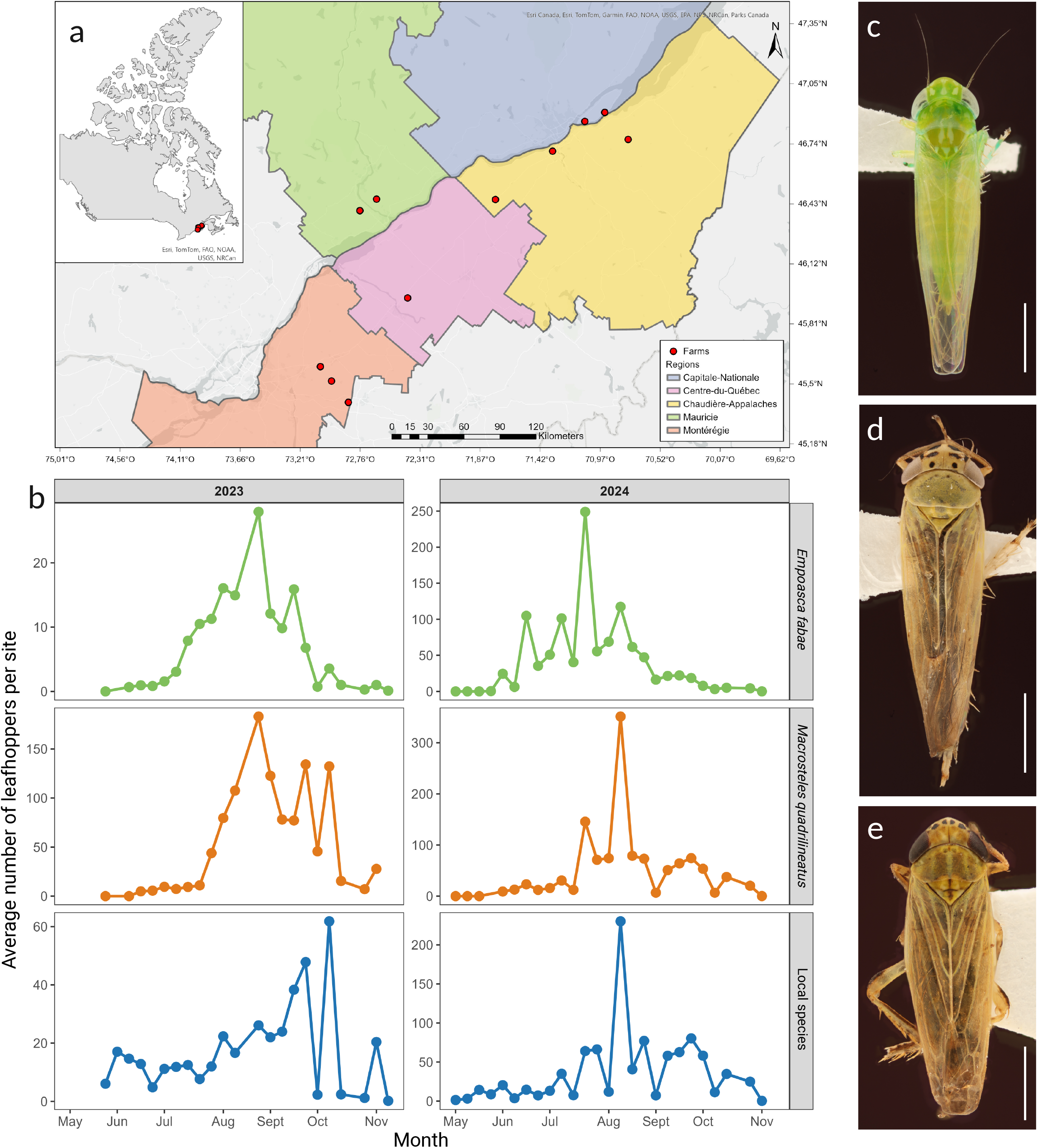
Leafhopper abundance and distribution across sampled strawberry fields. (**a**) Map showing the geographic locations of strawberry fields monitored in this study. (**b**) Seasonal dynamics of leafhopper abundance from May to November across both growing seasons (2023 and 2024), represented as the average number of leafhoppers per trap per site for each month. No error bars or standard deviations are shown, as the goal is simply to illustrate leafhoppers temporal dynamics. Color coding indicates taxonomic groups: green for *Empoasca fabae* (detailed in panel **c**), orange for *Macrosteles quadrilineatus* (detailed in panel panel **d**), and blue for local leafhopper species, with *Graminella nigrifrons* shown as an example (detailed in panel panel **e**). Scale bars in panels (**c**) to (**e**) represent 1 mm.

The sampling sites included strawberry fields either newly established or in their first or second year of fruit production, covering over 20 strawberry varieties (**Table S1**). Both short-day and day-neutral strawberry varieties were surveyed to capture the full range of strawberry field production systems in Québec. Leafhoppers were captured weekly from early May to late October using yellow sticky traps (Bug-Scan Dry Yellow, 10 cm X 25 cm, Plant Products) replaced weekly. At each site, five traps were installed along the central row of the field, evenly spaced to span its entire length. Traps were collected regularly and stored at 4⍰°C until leafhopper identification. Over 8,000 yellow sticky traps were processed between both growing seasons

To analyze the impact of weather on leafhopper abundance and parasitism, we collected daily temperature and precipitation data from each farm using the NASA Prediction of Worldwide Energy Resources (POWER) database (NASA 2024; version 2.4.3). We chose this database because (*i*) local stations were located far from the farms (min = 4.67 km, max = 16.71 km, average = 9.95 km), and (*ii*) for five farms, the stations did not have complete daily datasets for two months.

### Leafhopper identification and parasitism

Leafhoppers were removed from the sticky traps using an adhesive remover made from d-Limonene and sweet orange extract (Goo Gone, USA). Among the collected leafhoppers, only four species were identified to the species level: *E. fabae* and *M. quadrilineatus*, the two migratory and most abundant species in 2021 and 2022 (Plante et al. 2024), as well as *Japananus hyalinus* and *Dalbulus maidis*, newly recorded in Québec and Canada, respectively. All remaining species, hereafter referred to as “locals,” were identified to the genus level based on morphological characters, using taxonomic keys and identification guides (Plante et al. 2024). When identification was hindered by damage or color fading, male genitalia were dissected to achieve genus-level resolution. In such cases, sclerotized and fatty tissues were cleared by placing the genitalia in a 10% KOH tissue lysis buffer solution for 24 hours at 56⍰°C.

Additionally, prompted by our observations during the 2021–2022 sampling seasons of ectoparasitic larval sacs attached to leafhopper abdomens, presumably from the family Dryinidae (Plante et al. 2024, author observations), we recorded the number of parasitized leafhoppers per species, along with collection dates and locations, during the 2023–2024 sampling seasons. The presence of larval sacs (known as thylacium) (Virla et al. 2023), served as an indicator to estimate parasitism rates across sites and years.

### Statistical analyses Overview

All statistical analyses and graphic design were conducted using R version 4.4.2 (R Core Team, 2024) and RStudio version 2024.12.1+563, along with the following packages: DHARMa version 0.4.7 (Hartig 2024), ggplot2 version 3.5.1 (Wickham 2016), glmmTMB version 1.1.10 (Brooks et al. 2017), iNEXT version 3.0.1 (Chao et al. 2014; Hsieh and Chao, 2016), performance version 0.13.0 (Lüdecke et al. 2021), splines version 4.4.2 (R Core Team, 2024), and vegan version 2.6-10 (Oksanen et al. 2025).

### Leafhoppers diversity

Leafhopper diversity was compared between this study (2023–2024) and our previous survey (Plante et al. 2024) to evaluate whether diversity metrics differed across years. Because not all specimens in the current study were identified to species level, analyses were conducted at the genus level. Given differences in sampling effort, due to variations in the number of farms, traps, and trap deployment times, diversity was estimated using sample coverage-based rarefaction and extrapolation curves with Hill numbers (Chao and Jost, 2012; Roswell et al. 2021). This method provides more robust estimates than approaches such as equal-effort sampling or traditional rarefaction (Roswell et al. 2021).

Hill numbers integrate species richness and relative abundances into a unified framework across three orders (q values). When q = 0, all species are weighted equally, representing species richness. At q = 1, species are weighted in proportion to their relative abundance (analogous to the Shannon index). In contrast, at q = 2, greater weight is given to dominant species (analogous to the Simpson index) (Hill, 1973). Sample coverage exceeded 0.99 for each year, indicating high sampling completeness. For clear cross-year comparisons, we report both observed diversity and asymptotic estimates of total diversity for each q value.

### Seasonal leafhopper abundance

To assess whether the seasonal abundance (y-axis) of the three leafhopper groups (*E. fabae, M. quadrilineatus*, and local species) differed over time, we fitted separate generalized linear mixed models (GLMMs) for each group using the glmmTMB package. Modeling the groups individually was necessary, as including all three in a single model resulted in excessive multicollinearity (VIF > 42). In each model, Julian date (day of the year) was included as a fixed effect (x-axis), while farm (n = 11) and year (2023 and 2024) were treated as random effects. To account for variability in the number of traps per farm (due to some farms having two sites) and differences in trap deployment duration (in days), these factors were incorporated as offsets. To capture non-linear seasonal trends, we tested three regression splines (ns = 2, 3, and 4), using a negative binomial error distribution in all cases.

For each group, the best-fitting model was selected by simultaneously comparing the three candidate models using the performance package. This approach ranks models based on multiple fit metrics, including conditional and marginal R-squared, along with selection criteria such as AIC, AICc, and BIC, yielding an overall performance score from 0 to 100%, with higher scores indicating better model fit. Finally, for each group’s best model, we evaluated residuals for dispersion, zero-inflation, uniformity, and spatial and temporal autocorrelation using the DHARMa package (**Table 1, Table S2**).

**Table 1.**
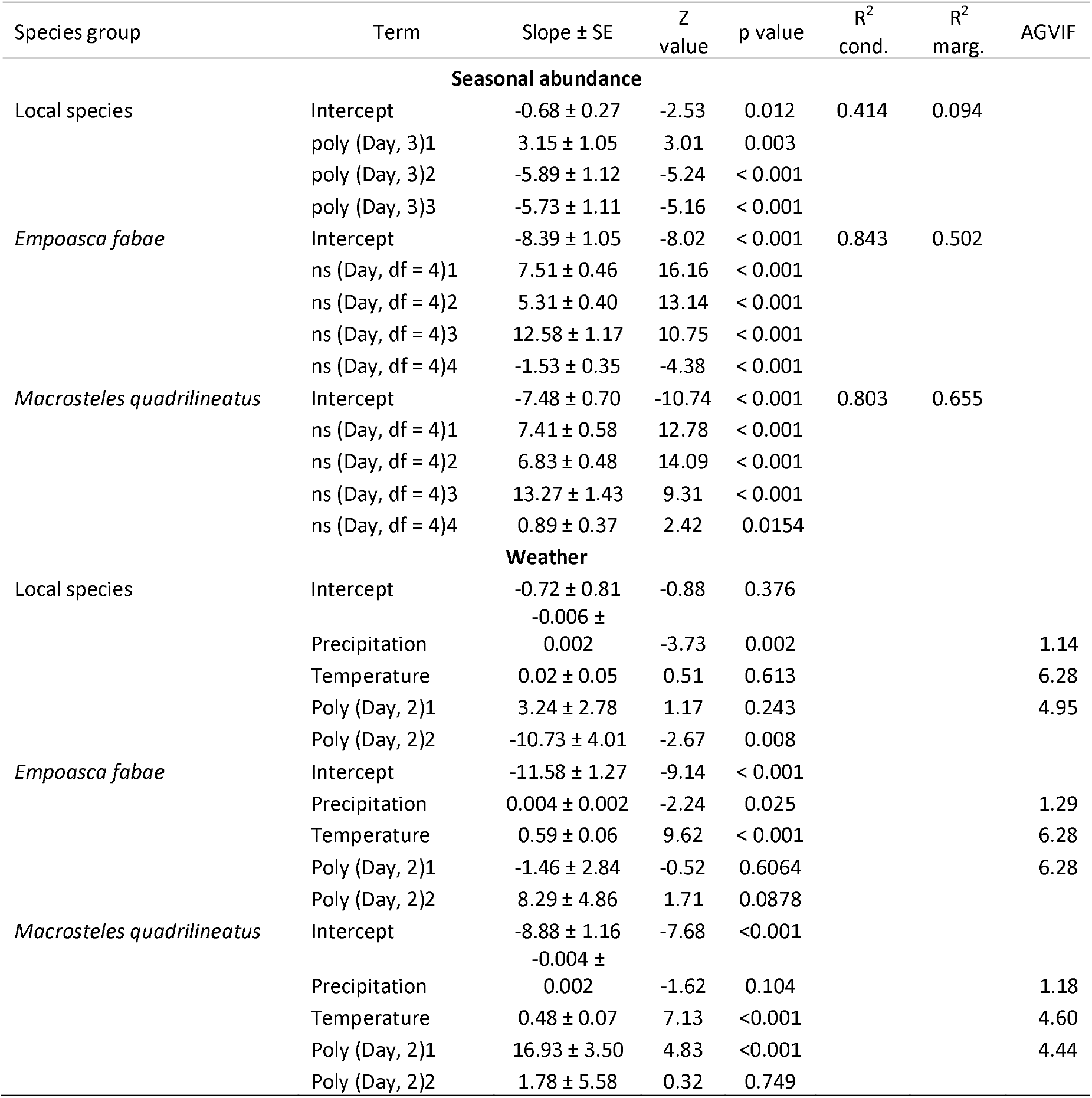
Summary of the generalized linear mixed models (GLMMs with negative binomial distribution) for local leafhopper species, *Empoasca fabae*, and *Macrosteles quadrilineatus* seasonal abundance (Julian date), as well as a function of temperature and precipitation averaged 30 days before trap removal in strawberry fields in Québec, Canada. Full details can be found in **Tables S2** and **S3**.

### Effect of climate on leafhopper abundance

The relationship between temperature, precipitation (x-axis), and leafhopper abundance (y-axis) was assessed using separate GLMMs for each species group, following the rationale described above. For each group, four models were fitted, incorporating lag times of 0, 7, 15, and 30 days for average temperature and accumulated precipitation prior to trap removal. Fixed effects included average temperature, accumulated precipitation, and Julian day (the latter modeled as a polynomial term to capture seasonal patterns) (**Table S3**). The year was included as a random effect. To address significant temporal autocorrelation identified in preliminary analyses, an autocorrelation structure accounting for repeated measures within farms was also incorporated. All models employed a negative binomial error distribution. Because the inclusion of an autocorrelation structure precluded the use of the performance package for model comparison, the best-fitting model for each species group was selected based on the lowest AIC and BIC values. Residual diagnostics for the selected models were conducted using the DHARMa package, evaluating dispersion, zero-inflation, uniformity, and spatial and temporal autocorrelation (**Table 1, Table S3**). Additionally, multicollinearity among predictors was assessed using the performance package.

### Insecticide application and leafhopper abundance

We investigated whether the abundance of leafhoppers (y-axis) changed before and after insecticide application for each species group (x-axis). To do so, we first collected information on the active ingredient and application date of insecticides at each site or farm. We then recorded the number of insects captured in traps both before and after the application date (**Table S4**). To ensure balanced comparisons, we normalized the number of leafhoppers captured per species group based on the number of days each trap remained active at a given site. Finally, we compared the normalized leafhopper counts before and after each insecticide application using a Wilcoxon paired test. This analysis was performed only for active ingredients applied at least four times.

### Parasitism occurrence, rates, and association with climatic parameters

To investigate whether parasitism occurrence varied seasonally and in relation to climatic parameters (model *i*), as well as to assess monthly changes in parasitism rates (model *ii*), we employed generalized linear mixed models (GLMMs). However, due to the low occurrence of parasitism within groups (see Results), we first analyzed seasonal parasitism occurrence as a binomial event (0 = no parasitism, 1 = parasitism) and pooled parasitism rates across the three leafhopper groups by month for subsequent analyses.

For model (*i*), we used a GLMM with a binomial error structure to test whether the probability of parasitism occurrence (y-axis) varied by month and was associated with climatic parameters (x-axis), specifically average temperature and accumulated precipitation. As in the leafhopper abundance models, we evaluated four-time lags (0, 7, 15, and 30 days) preceding trap removal. Fixed effects included Julian day (to capture seasonal effects), average temperature, and accumulated precipitation; offsets accounted for the total number of traps per farm and their deployment duration. Farm and year were included as random effects. The best-performing model across the four lag structures was identified using the performance package, based on multiple fit metrics and overall ranking. Residuals of the selected model were then assessed for dispersion, zero-inflation, uniformity, and spatial and temporal autocorrelation using DHARMa, along with checks for multicollinearity among predictors (**Table 1, Table S3**).

For model (*ii*), we examined whether monthly parasitism rates (y-axis) varied over the season (x-axis). Parasitism rates were aggregated across trap removal dates within each month, with month treated as a numerical variable (e.g., May = 5, June = 6). Year and farm were included as random effects, and models were fitted with a beta-binomial error distribution to model proportional data appropriately. To capture non-linear seasonal trends, we tested three natural spline transformations (ns = 2, 3, and 4) for the month effect. Models also included offsets for total traps and exposure duration, with total leafhopper abundance incorporated as a weight. Due to zero-inflation issues detected in preliminary models, a zero-inflation structure (ziformula = ~1) was specified. Model selection followed the same multi-metric approach using performance, and residual diagnostics were performed as above (**Table 1, Table S5**).

### Dryinidae capture and identification

Adult Dryinidae parasitoids were sampled from July to September 2024 on farms that exhibited the highest parasitism rates during the 2023 season, specifically in Montérégie (**Table S6**). Sampling was performed using a D-Vac insect aspirator fitted with a net, which was swept along the edges of strawberry fields and adjacent vegetation. This approach enabled the collection of parasitoids without disturbing insect populations within the fields, thereby avoiding interference with parasitism rate estimates. When Dryinidae individuals were observed in the net, a mouth aspirator was used to selectively extract them from the general catch. Additionally, one adult female Dryinidae specimen was collected from a yellow sticky trap in September 2024 (**Table S6**).

In total, 11 adult Dryinidae specimens were collected, all females, identified by their distinctive chelate protarsus (pincer-like forelegs), a key morphological feature of this family, commonly known as “pincer wasps”. To facilitate species-level identification, the chelae (pincers) of all specimens were slide-mounted, focusing on diagnostic characters of the protarsus such as the arolium, enlarged claw, and lamellae on the fifth protarsomere. For slide preparation, one foreleg tarsus from each specimen was detached and placed in boiling water for five minutes, then transferred to a 10% KOH solution and heated in a bain-marie (without boiling) for approximately five minutes. The tarsi were subsequently rinsed in 30% ethanol, cleaned, and transferred to lactic acid. Dehydration was achieved by placing them in 100% isopropanol overnight, with a glass slide gently pressed on top to maintain orientation for mounting. Tarsi were slide-mounted in Euparal mounting medium, whereas the remaining bodies were point-mounted for morphological examination. Specimens were identified using the taxonomic key of Olmi (1984) and by comparison with reference material from the Canadian National Collection of Insects, Arachnids, and Nematodes (hereafter CNC) (Agriculture and Agri-Food Canada, Ottawa, Ontario, Canada).

### Parasitoid DNA extraction and molecular identification

To comprehensively characterize the dryinid parasitoids on leafhoppers in Eastern Canada, we employed three complementary strategies:

#### (i) Parasitoid mitogenome reconstruction from parasitized M. quadrilineatus

Given that nearly all parasitized leafhoppers in our study were *M. quadrilineatus*, a species highly abundant in Québec’s strawberry fields (Plante et al. 2024), we focused our identification efforts on the parasitoids of this species. This approach was further justified by the availability of the *M. quadrilineatus* genome (Vasquez et al. 2023), which allowed us to filter out host reads during bioinformatic analyses. To capture the diversity of parasitism across our study area, parasitized *M. quadrilineatus*, identified by the presence of external larval sacs, were grouped by farm within each region. Specifically, we created three pooled subsamples, each containing 10 parasitized individuals from a single farm. Two subsamples came from two different farms in Montérégie, and the third from a farm in Centre-du-Québec (**Table S7**). Together, these sites represent a geographic gradient spanning our sampling area from the southernmost to the northernmost point. Collections took place between July and October 2023.

Total DNA was extracted following the protocol of Molligan et al. (2025) with minor modifications. Specimens were rinsed three times with sterile ddH_2_O, then parasitized leafhoppers were homogenized using a mini-pestle in 400 µL of prewarmed at 65⍰°C lysis buffer containing 20CmM EDTA (pH⍰8.0), 100CmM Tris-HCl (pH⍰7.5), 1.4⍰M NaCl, 2% (w/v) CTAB, and 4% (w/v) PVP-40. An additional 300⍰µL of the same buffer was added, and samples were incubated at 65⍰°C for 30 minutes with shaking at 300⍰rpm, inverting tubes every 10 minutes. The supernatant was extracted twice with 700⍰µL of chloroform–isoamyl alcohol (24:1). DNA was then precipitated using 70% ice-cold isopropanol, washed with 70% ethanol, air-dried for ~10 minutes, and finally eluted in 10⍰mM Tris-HCl, 0.1⍰mM EDTA (pH⍰8.0).

Library preparation and Illumina NovaSeq PE150bp paired-end sequencing were performed by the Genome Québec Innovation Centre (Montréal, Canada). Raw reads were quality filtered using BBTools (Bushnell, 2014) to remove sequencing adapters and low-quality bases. Host-derived reads were identified and removed by aligning the filtered data to the *Macrosteles quadrilineatus* reference genome (BioProject ID: PRJNA928792) using Bowtie2 (Langmead and Salzberg, 2012). Unaligned reads were retained for de novo assembly using SPAdes v4.0.0 in “careful” mode and stepped k-mer sizes up to 120 (Prjibelski et al. 2020). Contigs were filtered to retain those between 10–20 kb in length and with a coverage depth of 20–100×. Candidate mitochondrial contigs were identified by querying the NCBI nt database using BLASTn v2.13.0 (Altschul et al. 1990). A 16.5 kb contig with ~51× coverage was recovered from the Centre-du-Québec farm sample, showing high similarity to insect mitochondrial genomes. This contig was annotated using MITOS2 (Galaxy version 2.1.9⍰+ ⍰galaxy0) and visualized in Proksee (Bernt et al. 2013; Grant et al. 2023). All tRNA genes were validated using tRNAscan-SE v2.0 (Chan and Lowe, 2019), and intergenic regions between rRNA genes were manually reviewed. A GC-rich region was designated as the putative origin of replication.

#### (ii) Barcoding of parasitoid larva

A single parasitoid larva that had emerged from an external larval sac on the abdomen of *M. quadrilineatus*, collected on a yellow sticky trap, was selected for DNA extraction, as described above. For this larval sample, the *coxI* gene was amplified using LCO1490 and HCO2198 primers following the protocol recommended by Folmer et al. (1994). Sanger sequencing was performed at the Centre de recherche du CHU de Québec (Québec, Canada). Sequence alignment and cleaning were conducted in Geneious (Dotmatics, v.2024), and the resulting sequence was queried against the NCBI GenBank database using BLAST. Additionally, the larval *coxI* sequence was aligned with the mitochondrial genome constructed from the parasitized leafhoppers to confirm whether the identities matched.

#### (iii) Mitogenome reconstruction and barcoding of adult Dryinidae

##### DNA Extraction and Library Preparation

DNA extractions of the 11 Dryinidae adult specimens were performed using a DNeasy Blood and Tissue Kit (QIAGEN, Canada) following the procedure for a disposable microtube pestle as recommended by the manufacturer. In this case, each specimen tissue was homogenized individually. Library preparation and sequencing were carried out by Daicel Arbor BioSciences (Ann Arbor, MI, USA), which customized the protocols according to the preservation status of each sample using either ancient DNA (aDNA) or degraded DNA workflows, depending on the level of DNA fragmentation. Genomic DNA (gDNA) of the 11 adults and the larva previously used for PCR and Sanger sequencing were inspected for quality and quantity using spectrofluorimetric assays and TapeStation 4200 (Agilent). Most samples exhibited low molecular weight DNA profiles. Notably, specimen 3 showed extensive degradation and a DNA concentration below the required minimum threshold of 10 ng, so it was not further analyzed. Libraries were prepared using dual-indexed Illumina-compatible adapters, targeting an average insert size of ~300 bp, and sequenced on the Element Biosciences AVITI platform (PE150 model), with a target of ~10 Gbp per sample.

##### Read processing and genome assembly

Raw sequencing reads were cleaned using fastp v0.23.4, applying default quality filters (Phred ≥15) and adapter trimming. To ensure accurate trimming of potential reverse complement adapter sequences, the adapter_fasta option was used. Cleaned reads were assembled using SPAdes v4.1.0 in “careful” mode with multiple k-mers (21, 33, 55, 77, 99, 127). Contigs between 15,000-25,000 bp were extracted using awk and aligned to the NCBI nt database via BLASTn v1.16.0 to recover mitochondrial fragments. Hits aligning to *Gonatopus* were further filtered and annotated with MITOS2 using the invertebrate mitochondrial code and RefSeq89 Metazoa. Geneious Prime v2025.1.3 was used to adjust sequence orientation (trnQ as start), perform multiple sequence alignments, and estimate sequence identity. Synteny maps were generated with pyGenomeViz (Shimoyama 2024), using reciprocal BLAST.

##### Mitochondrial SNP calling and consensus building

Trimmed reads were mapped to the high-quality *Gonatopus* mitochondrial genome from the *Macrosteles* sample from Centre-du-Québec region using BWA-MEM v0.7.17 (Li 2013). BAM files were processed with SAMtools v1.8 (sorted, filtered for mapping quality ≥5, soft-clip <30 bp), and PCR duplicates were removed using Picard’s MarkDuplicates. Aligment around indels were refined with GATK v3.6 RealignerTargetCreator and IndelRealigner (DePristo et al. 2011). Overlapping paired-end reads were clipped using bamUtil v1.0.48 to avoid coverage inflation. Coverage statistics were calculated using samtools depth. Single-nucleotide polymorphisms (SNP) were called with BCFtools v1.15 (mpileup and call, using the multiallelic caller) using haploid settings and custom filtering (Danecek and McCarthy 2017). Genotypes supported by <4 reads or with low mapping/call quality were set to missing. To avoid artifacts from non-mitochondrial reads, a custom Python script filtered SNPs with ambiguous allele frequencies (0.1 < REF/(REF + ALT) < 0.9). Final consensus sequences were built using BCFtools consensus with masked missing or absent data.

##### Phylogenetic analysis

The *coxI* gene sequences from our samples were aligned with 17 *Gonatopus* COI sequences from NCBI (**Table S8**), the Sanger-sequenced larva *coxI*, and the *coxI* from the *Macrosteles*-associated *Gonatopus* and the wasp *Polistes dominula* as an outgroup. Sequences were aligned in Geneious using default parameters and automatic orientation detection. A phylogenetic tree was constructed using IQ-TREE with 1,000 ultrafast bootstraps and model selection. A second tree using concatenated genes (excluding those with >50% missing data in >3 samples) was generated, focusing on our samples only, using partitioned models. Both trees were visualized with iTOL and the COI tree was rooted at the midpoint.

##### Host prey and entomopathogen Identification

To infer the diversity of parasitoids hosts or prey, we downloaded all Cicadellidae *coxI* sequences from BOLD v3 (accessed 2025-07-12). A custom BLAST database was created and assembled contigs were queried using BLASTn v2.16.0. Hits with ≥90% identity and alignment length ≥50 bp were retained and parsed using awk. *Beauveria bassiana* sequences, a known entomopathogenic fungus, were unexpectedly detected in the dataset. To confirm its presence, BLASTn outputs from genome assemblies were screened to document its presence and potential role in parasitoid–host–microbe interactions.

## RESULTS

### Leafhopper abundance and diversity

Over the two sampling seasons, we captured a total of 82,130 leafhoppers: 30,246 in 2023 and 51,884 in 2024 (**Table S9**). In 2023, the abundance of *E. fabae* and *M. quadrilineatus* was low from May through mid-August, when both species reached a peak (**Fig.⍰1b**). Their numbers then declined steadily, approaching zero by the end of the season. Local leafhopper species were more numerous early in the season, peaking later in early October before also decreasing (**Fig.⍰1b**). In 2024, *E. fabae* exhibited a slightly earlier peak in late July, while *M. quadrilineatus* followed a pattern similar to 2023, peaking again in mid-August (**Fig.⍰1b**). Local species reached maximum abundance earlier than in the previous year, with a peak in early August (**Fig.⍰1b**). Despite these shifts in peak timing, the overall seasonal dynamics for the three groups were consistent across both years.

A total of 59 leafhopper genera were identified in 2023 and 64 genera in 2024, with a total of 69 general between both growing seasons (**Table S9**). The most abundant genera in 2023 were *Macrosteles, Empoasca, Graminella, Paraphlepsius*, and *Draeculacephala* (**Fig. 1c-e, Fig. 2a, Fig. S1**), whereas in 2024, *Hebata* spp. replaced *Draeculacephala* among the five most abundant (**Fig.⍰2a, Fig. S1**). Community composition also differed between years. In 2023, *Macrosteles* accounted for 68.47% of all captures, with *M. quadrilineatus* comprising 97.18% of that genus, and 66.53% of total leafhoppers (**Table S9**). *Empoasca* represented 8.75% of captures, entirely identified as *E. fabae* (**Table S9**). In 2024, the distribution was more balanced: *Macrosteles* made up 39.19%, with *M. quadrilineatus* accounting for 96.49% of that group, while *Empoasca* increased to 37.76%, again composed exclusively of *E. fabae* (**Table S9**).

**Figure 2.**
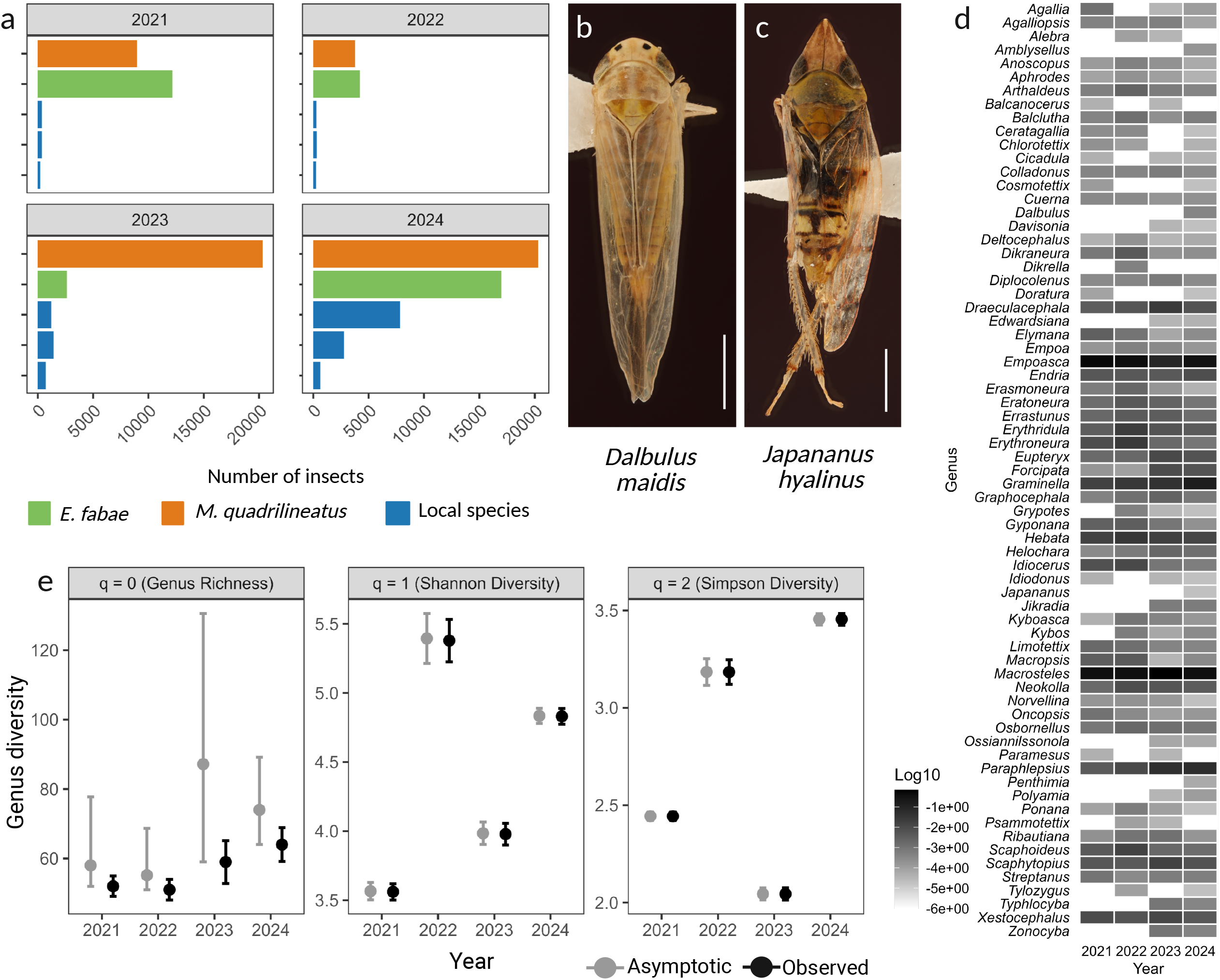
Leafhopper diversity recorded from 2021 to 2024. (**a**) The five most abundant genera detected in 2021 and 2022, compared with those identified in this study. *Empoasca fabae* (green) and *Macrosteles quadrilineatus* (orange) were consistently the most abundant, while the remaining genera were primarily local species (blue) (see Fig. **S1**). (**b**) Dalbulus maidis (corn leafhopper), reported here for the first time in Canada. (**c**) *Japananus hyalinus* (maple leafhopper), reported for the first time in Québec. Scale bars in panels (**b**) and (**c**) represent 1 mm. (**d**) Presence/absence and relative abundance of leafhopper genera detected from 2021 to 2024, highlighting the ‘core leafhopper diversity’ and distinguishing transient or less frequently detected local species. (**e**) Genus’s richness and alpha diversity comparisons (Shannon and Simpson indices) across growing seasons, including both observed and asymptotic estimates. Model estimates (black lines) and 95% confidence intervals (grey lines). A summary of the statistical analyses is presented in **Table S10.** Bar

Some leafhoppers genera were detected exclusively in a single year of the two sampled in this study. In 2023, we recorded *Alebra, Balcanocerus, Paramesus*, and *Psammotettix*, while nine genera were observed only in 2024: *Amblysellus, Ceratagallia, Chlorotettix, Cosmotettix, Dalbulus, Doratura, Japananus, Penthimia*, and *Tylozygus* (**Fig.⍰2b-d**). However, several of these genera were previously observed in 2021 and 2022 (**Fig. 2d**). Given the agricultural importance of *Dalbulus maidis*, the corn leafhopper, a major pest of corn in South and Central America, and more recently the United States, we identified all 20 *Dalbulus* specimens captured in 2024 in Québec to the species level. All were confirmed as *D. maidis* (**Fig.⍰2b, d, Table**⍰**S10**). These specimens were collected between September and October from five farms across the Montérégie, Mauricie, and Chaudière-Appalaches regions (**Table**⍰**S10**), marking the first documented record of this important pest in Canada. Due to the potentially significant economic impact of this species on corn production and the Canadian agricultural sector, a specimen was deposited in the CNC under voucher number CNC2151743, with Dr. Joel Kits overseeing the Hemiptera division. Similarly, a specimen belonging to the genus *Japananus* was identified as *Japananus hyalinus*, the maple leafhopper (**Fig.⍰2c, d, Table**⍰**S9**). Although this species has previously been reported in Canada, primarily in British Columbia and Ontario, and is occasionally noted in Québec via citizen science platforms like *iNaturalist*, this represents the first official record for Québec and the easternmost occurrence in Canada. Regarding the diversity metrics, genus’s richness (q = 0) did not vary significantly among years (**Fig. 2e, Table S11**). However, Hill Shannon diversity (q = 1) differed significantly across years, with the highest value observed in 2022, followed by 2024, 2023, and 2021. Similarly, the Hill Simpson diversity index (q = 2) also varied significantly, though with a different ranking: it was highest in 2024, followed by 2022, 2021, and lowest in 2023 (**Fig. 2e, Table S11**).

### Effect of Julian day and climatic variables on leafhopper abundance

Regarding the seasonal leafhopper abundance, the best-fitting models for all three leafhopper groups, *E. fabae, M. quadrilineatus*, and local species, revealed a significant influence of Julian date on seasonal abundance (*p*⍰< ⍰0.001) (**Table 1**). Fixed effects explained 50.2%, 65.5%, and 9.4% of the variance for *E. fabae, M. quadrilineatus*, and local species, respectively. No evidence of spatial or temporal autocorrelation was detected in residuals for any group, as indicated by Moran’s I and Durbin-Watson tests (**Table**⍰**S2**). Model curves showed distinct seasonal dynamics (**Fig.⍰3a, Table 1, Table S2**). For *E. fabae*, abundance remained low until early June, then increased steadily to peak in early July, followed by a gradual decline toward zero by late October to early November, aligning with the end of the crop season (**Fig.⍰3a, Table 1**). Similarly, *M. quadrilineatus* displayed an early-season trend but reached a more pronounced peak in early September before sharply declining (**Fig.⍰3a, Table 1**). In contrast, local species were already present in May, maintained relatively stable populations through August, and then slightly increased before tapering off by season’s end (**Fig.⍰3a, Table 1**).

**Figure 3.**
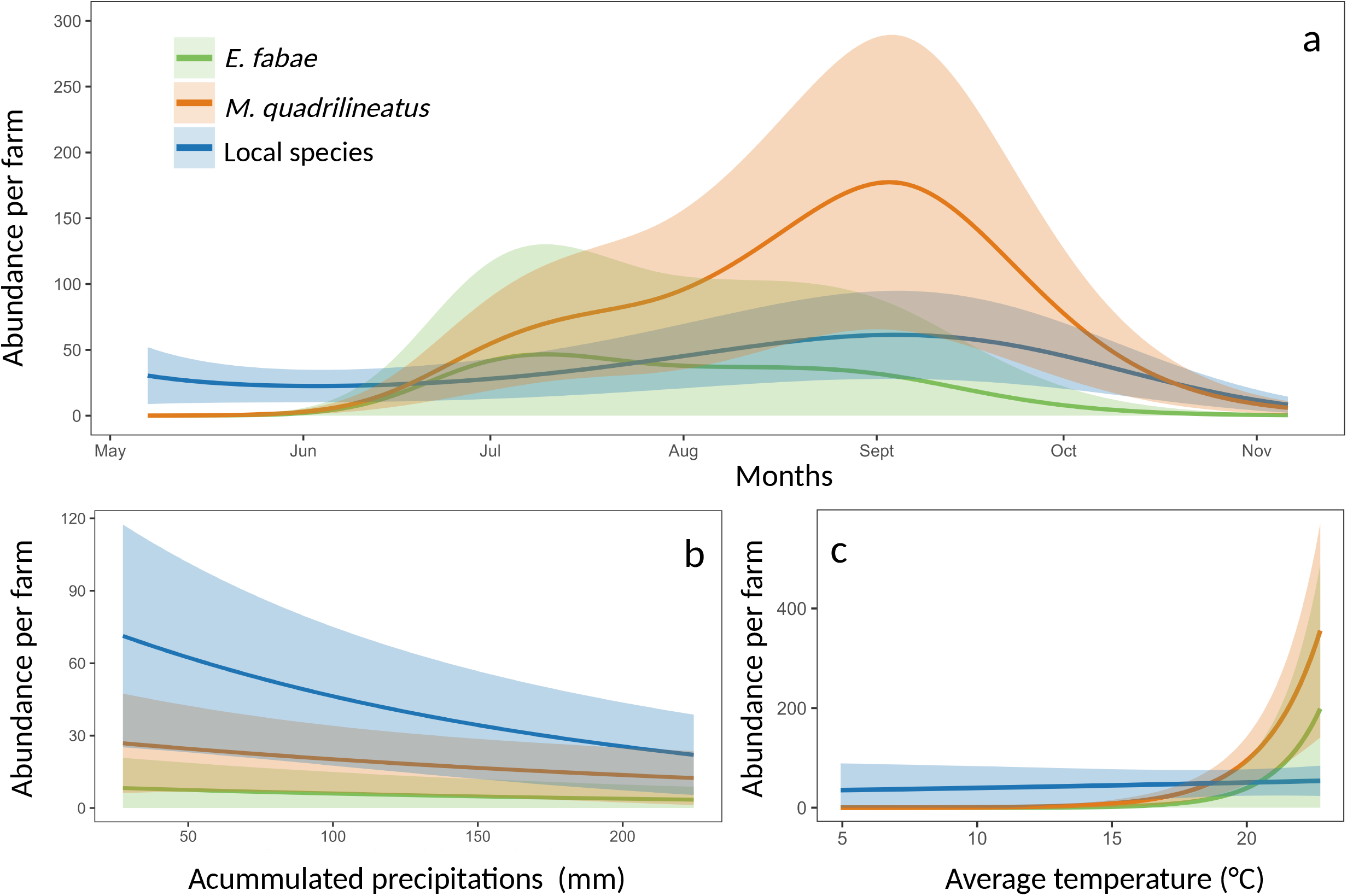
Leafhopper population structure in relation to temporal and climatic variables. **(a)** Seasonal variation in leafhopper population structure across the growing period. (**b**) Relationship between population structure and accumulated precipitation (mm). (**c**) Relationship between population structure and average temperature (°C). In all panels model estimated is presented by the dark color lines and the shadings represents the 95% confidence intervals. Corresponding statistical models are presented in **Table 1** and Supplementary **Tables S2** and **S3.** Color coding indicates taxonomic groups: green for Empoasca fabae, orange for Macrosteles quadrilineatus, and blue for local leafhopper species.

After testing four lag periods, the best-fitting climatic models for all three groups incorporated average temperature and accumulated precipitation over the 30 days before trap removal (**Table**⍰**S3**). Similarly to the previous models, no evidence of spatial or temporal autocorrelation was detected in residuals for any group, as indicated by Moran’s I and Durbin-Watson tests (**Table**⍰**S3**). For *E. fabae*, temperature had a significant positive effect on abundance (*p*C<C0.001) (**Fig.⍰3b, Table 1**), while precipitation had a significant negative effect (*p*C=C0.025) (**Fig.⍰3c, Table 1**). The model-predicted abundance remained low until the average temperature reached ~16⍰°C, after which it increased sharply (**Fig.⍰3a, Table 1**). Cumulative precipitation had little effect across the range of observed values (**Fig.⍰3c, Table 1**). For *M. quadrilineatus*, temperature had also a significant positive effect (*p*C<C0.001) (**Fig.⍰3b, Table 1)**, while precipitation was not significant (*p*C=C0.104) (**Fig.⍰3c, Table 1**). Abundance remained near zero until average temperatures exceeded ~14⍰°C, after which it rose steeply (**Fig.⍰3a, Table 1**). Although the model curve suggested a potential decrease with increasing precipitation, this was not statistically significant (**Fig.⍰3c, Table 1**). For local species, precipitation had a significant negative effect on abundance (pC=C0.002) (**Fig. 3c, Table 1**), while temperature showed no significant influence (*p*C=C0.613) (**Fig.⍰3b, Table 1**). The model predicted relatively constant but low abundance across temperature ranges, while abundance decreased with higher 30-day cumulative precipitation (**Fig.⍰3a-c, Table 1**). Together, these results highlight distinct climatic and seasonal drivers for the three groups. The abundance of *E. fabae* and *M. quadrilineatus* was primarily temperature-driven, with clear thresholds (~16⍰°C for *E. fabae* and ~14⍰°C for *M. quadrilineatus*) marking sharp increases in adult populations. In contrast, local species abundance appears to be primarily insensitive to temperature but negatively impacted by precipitation, with higher rainfall levels associated with reduced population sizes.

### Insecticide application and leafhopper group abundance

We obtained insecticide application data from five sites across three farms, revealing the use of a wide range of active ingredients (n = 13), with application frequencies ranging from 2 to 13 between both seasons (**Table S4**). Based on this, we assessed changes in the abundance of each leafhopper group for the nine active ingredients applied more than four times. Eight out of nine active ingredients did not appear to reduce leafhopper abundance (**Fig. 4, Fig. S2, Table S4**), including the three most frequently used insecticides, cypermethrin (n = 13), spinetoram (n = 12), and abamectin (n = 10) (**Fig. 4, Table S4**). Only applications of acetamiprid were associated with a significant decrease in the abundance of *E. fabae* (*p* = 0.031) and of local species (*p* = 0.036), but not of *M. quadrilineatus* (*p* = 0.078) (**Fig. S2**).

**Figure 4.**
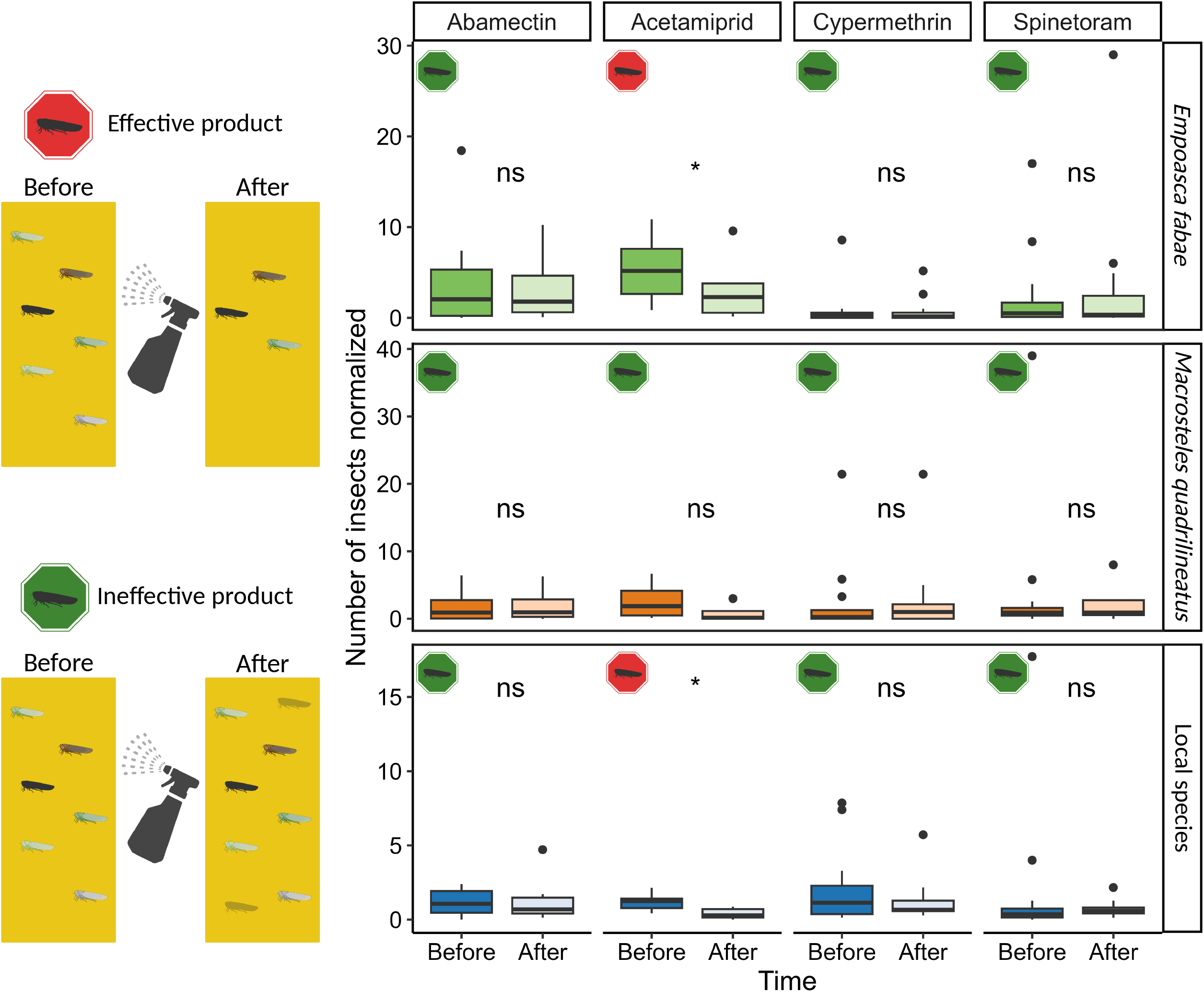
Effectiveness of insecticides on leafhopper population control. Impact of insecticides commonly used by strawberry growers on leafhopper abundance. Insecticides that significantly reduced leafhopper populations are marked with a red tag (effective), while those with no significant effect are marked with a green tag (ineffective). Among the nine insecticides applied more than four times between 2023 and 2024, we highlight the three most frequently used: cypermethrin (n = 13), spinetoram (n = 12), and abamectin (n = 10). Effects were evaluated across three leafhopper groups: the migratory species *Empoasca fabae* and *Macrosteles quadrilineatus*, as well as local species. Only acetamiprid showed a significant reduction in the abundance of *E. fabae* and local species. The remaining insecticides, including the ones shown here, did not lead to significant population decreases. Results for the other insecticides are provided in **Figure S2** and **Table S4**.

### Parasitism rates and the effect of climatic variables on parasitism

Because parasitism rates were low in certain months (**Table S12**), data from both sampling years were combined for analysis. Apparent differences emerged among *E. fabae, M. quadrilineatus*, and locals (**Table S12**). *Empoasca fabae* exhibited very low parasitism, with parasitized individuals detected only in July, August, and September, averaging about three specimens per month (**Fig.⍰5a-b, Table S12**). The proportion of parasitized *E. fabae* relative to total parasitized leafhoppers per month ranged from 0% to 1% (**Fig.⍰5a**). In contrast, *M. quadrilineatus* (**Fig. S3a**) had the highest parasitism rates, with parasitized individuals present from May through October (**Fig.⍰5a**). This species accounted for 78% to 97% of all parasitized leafhoppers monthly, peaking in July (**Fig.⍰5a-b**). The number of parasitized *M. quadrilineatus* ranged from 4 to 841 per month, with the highest count in September (**Fig.⍰5a**). Local leafhopper species were parasitized throughout the season as well, with monthly counts from 1 to 32 individuals (**Fig.⍰5a**). August and September had the highest numbers, while local species represented 4% to 22% of total parasitized specimens, peaking in October (**Fig.⍰5b**). Among these, *Graminella* spp. (**Fig. S3b**) were most frequently parasitized, followed by *Forcipata* spp., and equally parasitized *Kyboasca, Scaphytopius*, and *Zonocyba* spp. (**Table S12**).

**Figure 5.**
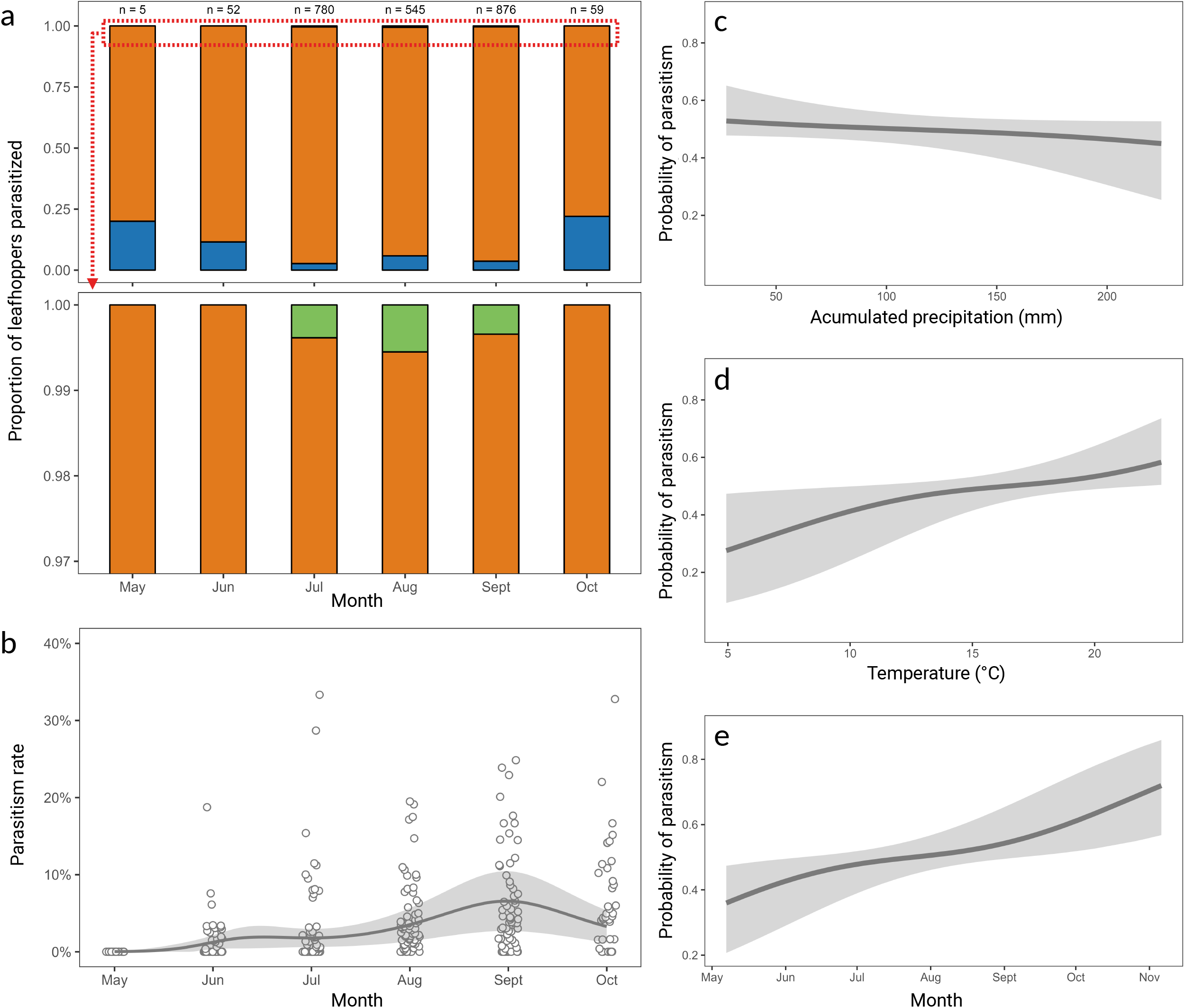
Parasitism rate and structure in relation to temporal and climatic variables. (**a**) Proportion of parasitized leafhoppers per month in 2023 and 2024. Data from both years were combined due to low overall parasitism rates. The bottom panel shows a magnified view (0.97– 1.00) highlighting *Empoasca fabae* (green), with parasitism primarily detected from July to September. (**b**) Monthly parasitism rates, averaging around 3% but occasionally exceeding 30% in isolated cases. (**c**) Relationship between parasitism and accumulated precipitation (mm). (**d**) Relationship between parasitism and average temperature (°C). (**e**) Seasonal variation in parasitism across the growing period. From c to e, model estimated is presented by the dark grey lines and the shadings represents the 95% confidence intervals. Statistical models corresponding to these results are presented in **Table 2** and Supplementary **Table S5.** Color coding indicates taxonomic groups: green for *Empoasca fabae*, orange for *Macrosteles quadrilineatus*, and blue for local leafhopper species.

Testing four different climatic models identified the best-fitting model as including average temperature and cumulative precipitation over a 30-day lag (**Table 2, Table S5**). Multicollinearity was low (VIF = 1.95 for both temperature and precipitation, 1.04 for Julian day) (**Table 2, Table S5**). Precipitation had no significant effect on parasitism occurrence (*p* = 0.103) (**Fig. 5c, Table 2**), while temperature showed a significant positive effect (*p* = 0.001) (**Fig. 5d, Table 2**). Julian day also had a significant positive effect (*p* < 0.001) (**Fig. 5e, Table 2**). The model’s marginal R^2^ indicated that fixed effects explained 24.8% of the variance (**Table S5**). Residual diagnostics found no spatial (*p* = 0.9332) or temporal (*p* = 0.9162) autocorrelation (**Table S5**). Predictions showed that the probability of observing parasitism remained stable around 50% across the range of cumulative precipitation values (**Fig. 5c, Table 2**). However, the probability increased markedly with temperature, rising from about 30% at a 30-day average of 5°C to nearly 60% at 22.5°C (**Fig. 4d, Table 1**). Similarly, predicted probabilities increased over the season, from roughly 35% in early May to above 70% by early November (**Fig. 5e, Table 2**).

**Table 2.**
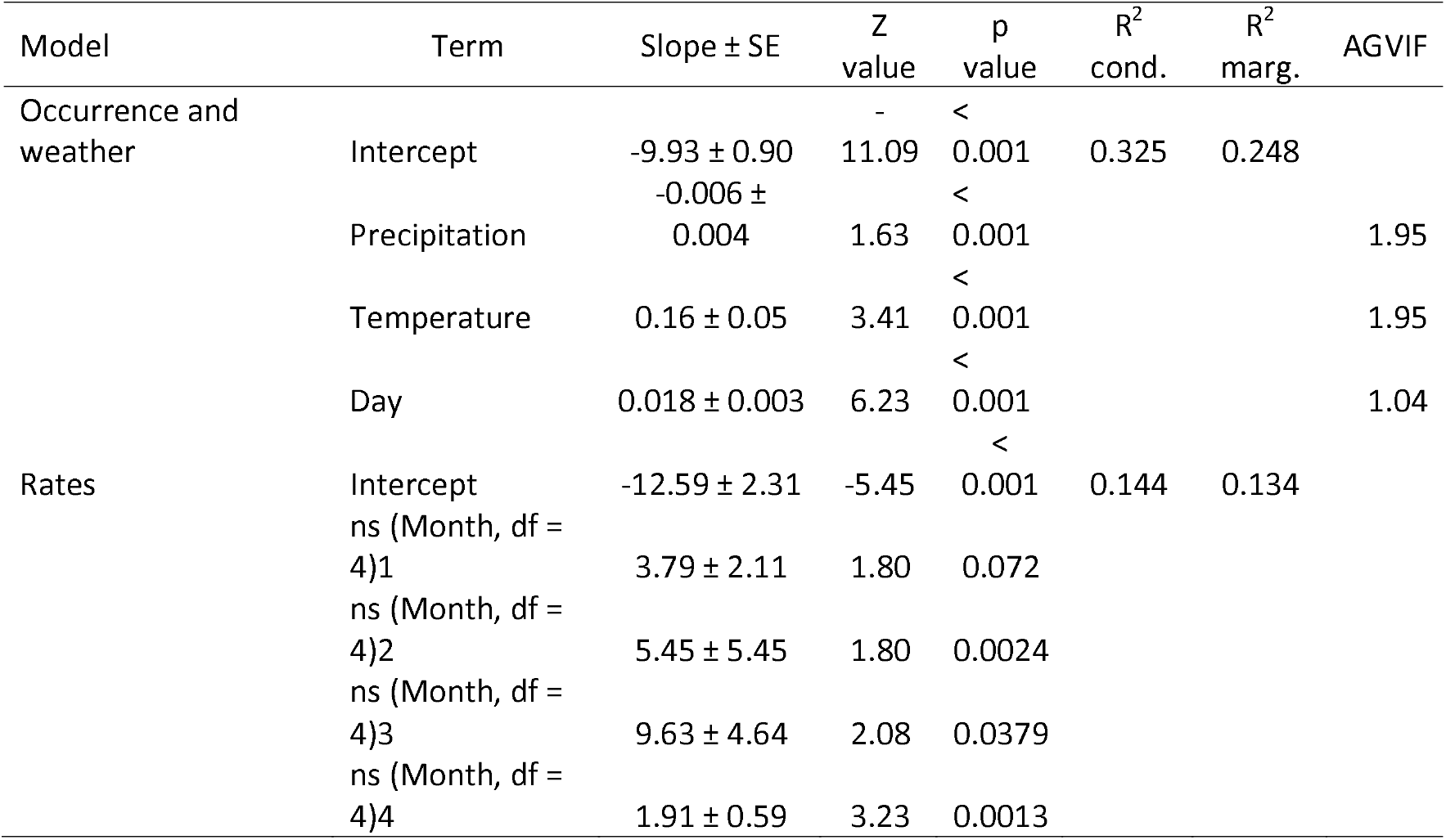
Summary of the generalized linear mixed models (GLMMs with binomial and beta-binomial distribution, respectively) for parasitism occurrence, weather (precipitation and temperature), and rates across months in strawberry fields in Québec, Canada. Full details can be found in **Table S4**.

Parasitism rates varied by month (*p* = 0.001), with a marginal R^2^ of 13.4%. No spatial (*p* = 0.2046) or temporal (*p* = 0.5318) autocorrelation was detected (**Table S5**). Individual samples ranged from 0% to 33% of parasitism, with all months containing at least some samples showing no parasitism (**Fig.⍰5b)**. The highest individual rates were observed in July and October (**Fig.⍰5b)**. Mean monthly parasitism rates ranged from 0% in May to nearly 6% in October, with an overall average of ~3%. Model predictions indicated parasitism rates near zero early in the season, followed by a gradual increase through the summer, a slight peak between June and July, and the highest rates (6–7%) in September before a decline in October (**Fig.⍰5b)**. Together, these results points that parasitism in Eastern Canadian agroecosystems is primarily driven by temperature and seasonal progression, with higher temperatures and advancing season significantly increasing the likelihood and rates of parasitism, while precipitation plays a lesser role.

### Parasitoids taxonomic identification

Using the taxonomic keys and diagnostic characters provided by Olmi (1984), we identified our 11 specimens from Québec as belonging to the genus *Gonatopus* (**Fig. 6a-c**). We further employed the same reference and the updated Nearctic *Gonatopus* key by Olmi (1993) to attempt species-level identification, followed by a comparative analysis with the more recent work of Guglielmino et al. (2018), which expanded the knowledge of Nearctic *Gonatopus* diversity. We also compared our material with authenticated specimens of 27 different *Gonatopus* species present at the CNC. Because the labial palpi of our specimens were either damaged or difficult to observe, we tested both branches of the dichotomous key concerning palpal segmentation (in one path, the labial palpi were considered to have two segments, in the other, three).

**Figure 6.**
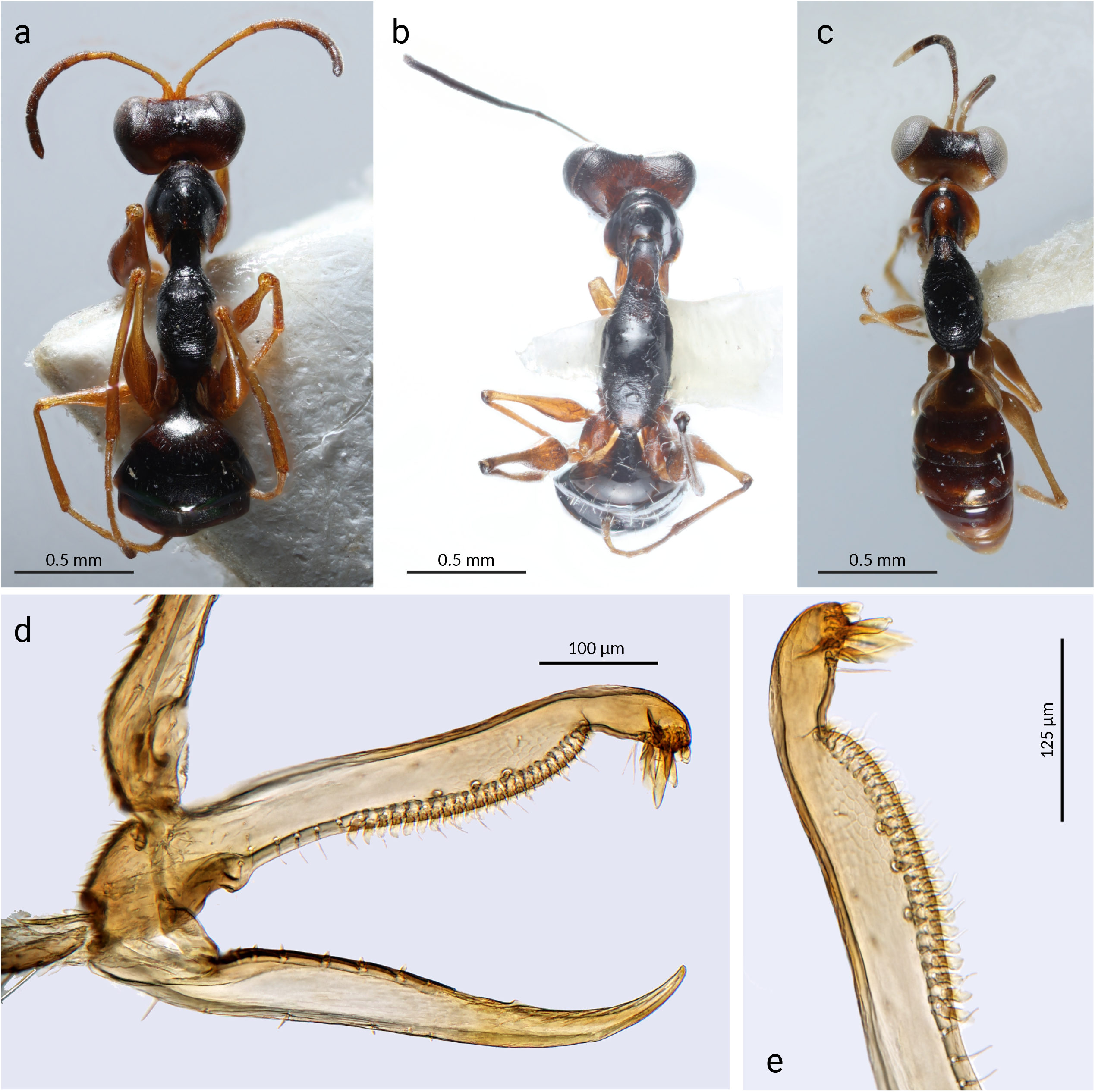
Morphologic representation of *Gonatopus* species. (**a, c**) Dorsal views of *Gonatopus contortulus* and *Gonatopus stenocrani*, respectively, from the Canadian National Collection of Insects, Arachnids, and Nematodes. (**b**) Dorsal view of *Gonatopus* sp., representing nine of the eleven adult specimens collected in this study. This lineage is putatively a new species for Canada. (d, e) Close-up views of the chelate protarsus of the Specimen 8 *Gonatopus* sp., showing two rows of lamellae (3 + 24).

Following the identification path for specimens with three-segmented labial palpi, we reached a dichotomy between two species, *Gonatopus freytagi* and *Gonatopus rapax*, yet neither fully matched our specimens. Our specimens lacked lateral points on the scutum and presented two rows of long lamellae on segment V of the fore tarsus. While *G. freytagi* shares the absence of scutum lateral points, it is described as having two rows of short lamellae. Conversely, *G. rapax*, corresponding to the alternative dichotomic choice, exhibits a scutum with two lateral points and two rows of long lamellae. Thus, our specimens partially resembled both species but did not correspond entirely to either. Considering the importance of chelate protarsi in the identification of female *Gonatopus*, we further compared the morphology of protarsomere V in our specimens with those of *G. freytagi* and *G. rapax*. The observed traits did not align well with either. *Gonatopus* group of 9–15 lamellae. On the other hand, *G. rapax* presents two peg-like hairs, two rows of 3+21 lamellae, and an apical group of at least 20. In contrast, our specimens exhibited three to nine peg-like hairs, 0–5 + 16–24 long lamellae, and an apical group of only 5–9 lamellae (**Fig. 6d–e**). Specimen 11 appeared to have a single row of lamellae on protarsomere V, but due to damage from collection on a yellow sticky trap, the exact count could not be determined (**Fig. S4**). When following the key for specimens with two-segmented labial palpi, Specimens 1, 3, 4, 5, 6, 7, 9, 10, and 11 keyed to *Gonatopus ashmeadi*, based on their lamellae being roughly equal in length. Meanwhile, Specimens 2, 5, and 8 were identified as *Gonatopus ombrodes*, given that their lamellae appeared longer near the base. However, closer morphological examination again revealed inconsistencies. *Gonatopus ashmeadi* is characterized by a single row of 8–20 lamellae of nearly equal length and an apical group of 6–8 lamellae, including 2–3 very long ones. Among our specimens, only Specimen 7 exhibited a comparable configuration, with a single row of 16 lamellae and an apical group of 5–6 lamellae, which still did not fully match the description. Similarly, *G. ombrodes* is described as having a single row of 14 lamellae, with the longest near the proximal end, and an apical group of 8 lamellae, two of which are very long, traits not observed in our specimens.

To explore the possibility that our specimens belonged to *Gonatopus* species not yet recorded in the Nearctic, we explored and applied the Palaearctic and Neotropical keys for female *Gonatopus* species (Olmi 1984). In the Palaearctic key, Specimen 10 keyed out as *Gonatopus sepsoides* (now considered a synonym of *Gonatopus clavipes*). This specimen exhibited the following diagnostic characters: segment I of the fore tarsus shorter than segment IV; inner side of segment V proximally not serrate; metathorax + propodeum lacking a median furrow; meso-metapleural suture distinct and complete; anterior surface of the metathorax + propodeum smooth and unsculptured (**Fig. S5**). These traits matched the species description, and identification was further supported by the morphology of the fifth protarsus, which had a single row of 11–12 lamellae beginning on the proximal third of segment V and an apical group of 8–9 lamellae.

For the remaining specimens, neither the Palaearctic nor Neotropical keys yielded satisfactory identifications. Discrepancies were confirmed by comparing the actual morphology of protarsomere V with the descriptions of proposed species from the keys. Taken together, these morphological differences suggest the presence of at least two distinct *Gonatopus* species among the 11 specimens analyzed, including forms that do not correspond to any species currently described in the Nearctic, Palaearctic, or Neotropical keys.

### Molecular diversity of parasitoids and their hosts

To investigate the molecular diversity of parasitoids and their leafhopper hosts, we sequenced 10 Dryinidae adult specimens and one larva collected from parasitized *M. quadrilineatus*. Three pooled samples of parasitized *M. quadrilineatus* were sequenced separately. A complete mitochondrial genome of 16,502 bp was recovered from one pooled aster leafhoppers (~51× coverage) (**Fig. 7a, Table S13**). The genome exhibited the canonical metazoan mitochondrial architecture, including 13 protein-coding genes (PCGs), 24 tRNAs, and two rRNAs. As expected in insects (Molligan et al. 2025), dual tRNA families were observed for serine (*trnS1/tct, trnS2/tga*) and leucine (*trnL1/tag, trnL2/taa*) (**Fig. 7a**). Additionally, a Sanger-sequenced *coxI* fragment from a larva aligned perfectly (682 bp, 100% identity) with this mitogenome (**Fig. 7a**), confirming species identity as the same that is parasitizing *M. quadrilineatus* in the field.

**Figure 7.**
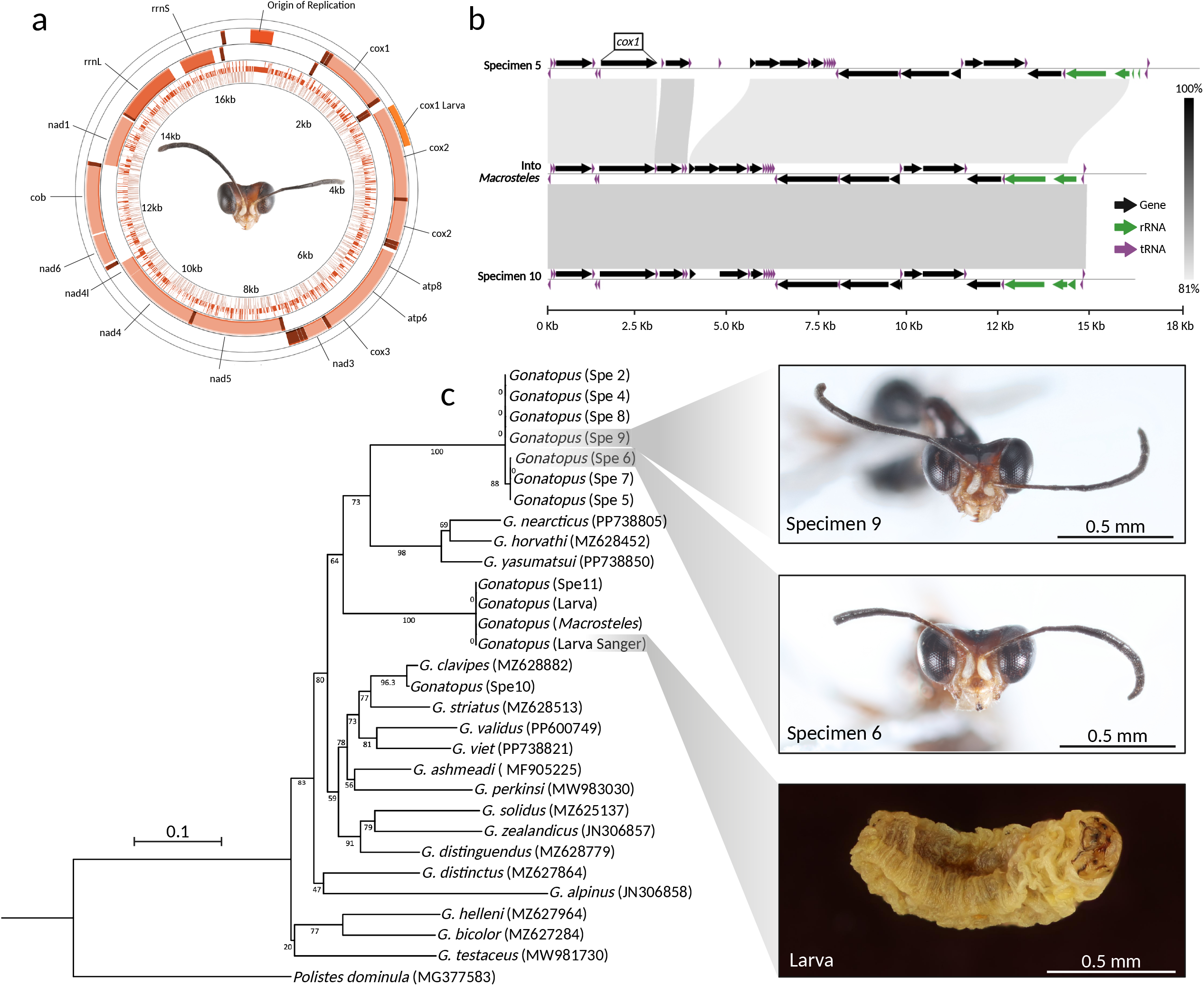
Molecular identification of *Gonatopus* specimens collected in this study. (**a**) Mitogenome of *Gonatopus* sp. reconstructed from *Macrosteles quadrilineatus* individuals parasitized by this wasp. The mitogenome includes key features such as the origin of replication and *cox* genes. A partial *cox1* gene from a larva (sequenced via Sanger) was found to be 100% identical to the corresponding region in the mitogenome. The orange color indicates that the host (*M. quadrilineatus*) was collected in Montérégie (see **Figure 1a**). Full mitogenome annotation is provided in **Table S13.** (**b**) Synteny analysis among three *Gonatopus* mitogenomes constructed in this study: one obtained from parasitized *M. quadrilineatus*, and two others from specimens 5 and 10. The results show that the *Gonatopus* sp. infecting *M. quadrilineatus* is more closely related to specimen 10 than to specimen 5. Annotations are available in **Table S15-S16**. (**c**) Maximum likelihood phylogenetic tree of *cox1* sequences for Gonatopus species available in GenBank (accession numbers in parentheses). Specimen 10 (see Figure S4) clusters closely with G. clavipes. Specimen 11 (Figure S5) groups with the sequence obtained from the parasitized Macrosteles individuals and with the cox1 sequence recovered from the larva using both Sanger and ancient DNA workflows. The remaining seven specimens form two distinct, yet related, clades. Bootstrap support was calculated with 1,000 replicates; the scale bar represents 1 substitution per 10 nucleotide positions. The European paper wasp *Polistes dominula* was used as the outgroup. Frontal views of specimens 6 and 9 are shown as examples of what may represent a cryptic novel taxon present in Eastern Canada. The corresponding larval specimen analyzed is also illustrated. A concatenated phylogenetic tree using the mitochondrial genes *atp6, atp8, cob, cox1, cox2, cox3, nad1, nad2, nad3, nad4, nad4l, nad5*, and *nad6* is presented in **Figure S6**.

For the 10 Dryinidae adult specimens and one larva, seven were processed under ancient DNA workflows and four under degraded DNA protocols. Fragment sizes were generally short (<1,500 bp), except for Specimen 1, which exhibited a broader length distribution. Sequencing yielded 49– 106 million reads per sample (median = 65 million), with filtered read lengths averaging 88–114 bp (**Table S14**). BLASTn searches identified 10 putative *de novo* assembled mitochondria across 6 samples, of which two show similarity to *Gonatopus clavipes* in Specimen 5 (86.6%) and Specimen 10 (99.9%), the later likely corresponding to the same species.

To further explore mitogenome structure and variation, synteny analyses were performed including the *M. quadrilineatus*-associated mitogenome. These analyses confirmed conserved gene content among parasitoid specimens but revealed notable structural variation in non-coding regions (**Fig. 7b**). For example, Specimen 10 contained a ~1,700 bp indel between *cox2* and *atp6*, resulting in a mitogenome approximately 3,000 bp longer than expected (**Table S15**). Specimen 5 exhibited a 986 bp insertion in the same region, whereas the insertion in Specimen 10 was 227 bp shorter. Given the read’s quality, at least part of these variants might originate from the miss-assembly of complex regions. Full-length alignments between the three specimens yielded only 74.2% sequence identity, a value likely inflated due to assembly artifacts such as rrnS fragmentation and gene duplications (e.g., *nad4L* duplication in Specimen 10 (**Table S16**). Despite these limitations, the results indicate the presence of at least three distinct *Gonatopus* species in the samples analyzed.

The partial *coxI* gene was recovered from all specimens except Specimen 1. A 545 bp alignment region shared across Dryinidae adult specimens, larva, and 18 NCBI *Gonatopus* sequences was used for phylogenetic inference. The resulting tree identified three well-supported clades (>95% bootstrap): one including Specimen 10, clustering with *G. clavipes* (with >99% identity) as supported by mitogenome nucleotide identity results; another with Specimen 11 and the larva (the sequences obtained using Sanger and AVITI), grouped with the *Macrosteles*-derived mitogenome (100% identity); and a third comprising the remaining specimens 2, and 4 to 9 (**Fig. 7c**). Clades 2 and 3 grouped closer to the *coxI* gene available in the NCBI for *G. nearcticus, G. horvathi*, and *G. yasumatsui*, although deep-node support was modest ranging from 45–75% (**Fig. 7c**). A second tree based on concatenated mitochondrial genes (*atp6, atp8, cob, cox1, cox2, cox3, nad1, nad2, nad3, nad4, nad4l, nad5*, and *nad6*) reproduced this topology, reinforcing the presence of at least three distinct species in Eastern Canada strawberry fields (**Fig. S6, Table S17**). Interestingly, this molecular detective work also provided insight into the feeding habits of these parasitoids, as they also act as leafhopper predators. A *coxI*-based host barcoding revealed that Specimen 7 had fed on *Macrosteles quadrilineatus* (100% identity), while Specimen 5 had consumed *Graminella nigrifrons* (98.1%) (**Table S18**). Specimen 4 showed partial matches to *Paraphlepsius* and *Psammotettix* species (92–93% identity over 61–78 bp). One partial *coxI* aligned to *Phlogotettix* with less than 80% identity (**Table S18**). However, since this genus is not found in Canada, has only been reported in Asia and Europe (Meshram et al. 2015), and because the reverse BLAST yielded no match, this alignment likely reflects a misidentification or an error in the NCBI reference data.

### Detection of Beauveria bassiana: uncovering a tri-trophic web

Amid the host-parasitoid dynamics, our assemblies revealed an unexpected third player: *B. bassiana*, a well-known entomopathogenic fungus. BLASTn analysis confirmed the presence of *B. bassiana* sequences in multiple genome assemblies (**Table S19**). While the origin remains unclear, potential explanations include environmental contamination, latent fungal infections in hosts or parasitoids, or incidental ingestion. However, we did not detect any *B. bassiana* reads in the *M. quadrilineatus* samples. Nonetheless, a mitogenome of this entomopathogenic fungus was reconstructed from specimen 6 (**Fig. S7**), although it was also detected in specimen 7 and specimen 10, which showed a white mycelium resembling that typically produced by the fungus (**Fig. S4**).

## DISCUSSION

This study provides the most comprehensive analysis to date of leafhopper dynamics and their parasitoid enemies in Canadian agroecosystems, offering ecological, genomic, and applied insights that are both novel and timely in the face of accelerating climate change. Across two consecutive growing seasons and over 8,000 sticky traps, we documented more than 82,000 leafhopper specimens identified to the genus level. Among these, migratory species such as *E. fabae* and *M. quadrilineatus* were the most abundant, as previously reported (Plante et al. 2024; Santos et al. 2025), exhibiting well-defined, temperature-driven phenologies. Although local species exhibited similar temporal distributions, with abundance peaks from mid-July to mid-August, they seemed more affected by precipitation, suggesting distinct ecological strategies between migratory and local taxa.

Migratory and local leafhopper species often exhibit distinct ecological and phenological traits, leading to different responses to factors such as climate, host plant availability, and natural enemies (Gatehouse 1997; Chapman et al. 2015). Understanding how these factors influence population dynamics is essential for pest species, especially under climatic conditions that may increase pest pressure (Nault and Ammar, 1989). This is critical for species involved in the transmission of plant pathogens (Plante et al. 2024; Santos et al. 2024; 2025). Effective population control necessitates a comprehensive understanding of leafhopper biology and seasonal dynamics to implement targeted and sustainable management strategies (Santos et al. 2025).

The potato leafhopper, *E. fabae* is known as a migratory species. It does not overwinter in Québec and it has been suggested to migrate at the end of the season to the extensive pine forests of the Gulf Coast States to overwinter (Taylor and Shields, 1995; Santos et al. 2025). Around late May and early June, as temperatures become more favorable, the species takes advantage of warm southerly air currents to move northward into its summer range (Taylor and Shields, 2018), including Eastern Canada (Plante et al. 2024; Santos et al. 2024; 2025). Given that nearly all *E. fabae* captured in yellow sticky traps are adults, the increase in trap catches observed in late June or July likely corresponds to the emergence of the first summer generation of adults, which developed from eggs laid by the migratory females (Santos et al. 2025). Similarly, the abundance peaks observed in late July or August may be the result of a second local generation (Santos et al. 2025). Since the average lifespan of this species ranges from 30 to 60 days, there is often generational overlap during this time of year (Decker and Cunningham, 1967; DeLong 1938; Hogg 1985; Hogg and Hoffman, 1989; Santos et al. 2025). Toward the end of the season, population abundance tends to decline, likely due to reduced host plant availability and cooler temperatures that are less suitable for leafhopper development. The migration of *E. fabae* relies on specific atmospheric conditions, including the presence of wind corridors located between 1,000 and 2,000 meters in altitude, at pressures of 800 to 900 hPa, and temperatures between 10 and 14⍰°C sustained over two to three days (Taylor and Reling, 1986, Taylor and Shields, 2018). These strict requirements limit the number of favorable migration opportunities. As a result, the timing of such conditions can vary from year to year, which may explain the observed differences in the timing of population increases between 2023 and 2024, thus shifting abundance patterns.

The aster leafhopper, *M. quadrilineatus* is a species that overwinters in northern regions of its range, such as southern Canada, by entering winter diapause in the egg stage (Westdal et al. 1961; Frost et al. 2013). The optimal temperature range for its survival and reproduction is between 5⍰°C and 20⍰°C (Bahar et al. 2018). In Québec, these temperature conditions are typically reached in May, when average monthly temperatures range from 10⍰°C to 15⍰°C (MELCCFP, 2025), likely marking the end of *M. quadrilineatus* diapause. Since the developmental period from egg to adult spans approximately 34 days (Hagel and Landis, 1967), the increase in adult captures observed in June likely corresponds to the emergence of individuals that overwintered in diapause. In addition to local emergence, the arrival of migratory individuals from the United States may also contribute to the rise in adult populations in June. Like *E. fabae, M. quadrilineatus* migrates in June from the southern United States into the American Midwest and Canadian Prairies using atmospheric currents and weather systems (Lee and Robinson, 1958; Nichiporick 1965; Wallis 1962). Given that leafhopper development typically takes a bit more than 30 days, both local and migratory individuals are likely to produce a first generation of locally emerging adults by July. The abundance peaks observed in August in both 2023 and 2024 are likely to result from overlapping generations (Hagel and Landis, 1967), specifically, the first and second local generations as well as early-season migrants.

The species considered local may overwinter in diapause as eggs, nymphs, or adults. Phenology of these species varies depending on overwintering strategy, but adults may be present from the beginning of the season because they have overwintered locally. Since no active migration should be involved, their population development does not rely on specific weather events, unlike migratory leafhoppers. Local species may benefit from early-season warming, which allows them to terminate diapause and initiate development earlier than migratory leafhoppers. This pattern is also observed in *M. quadrilineatus*, a species that, although not entirely local, partially overwinters *in situ*. This likely explains the higher abundance of this group at the start of both seasons studied here. Additionally, local species may benefit from the early availability of host plants, in contrast to migratory leafhoppers that potentially target more developed or specific hosts appearing later in the season. The fact that the population dynamics of *M. quadrilineatus* and the local species follow very similar patterns throughout the season supports the hypothesis that *M. quadrilineatus* overwinters in Québec.

One key limitation of the study is the grouping of several species under the category of “local species”, based on the assumption that they overwinter in Québec. However, most of these species have not been studied individually, particularly in their detailed biology and ecology. As a result, their overwintering strategy remains uncertain. It is therefore possible that some of the species included in the local group may exhibit migratory behavior. If so, this could influence the observed seasonal patterns attributed to local species and may have led to an overestimation of the early-season abundance attributed solely to local taxa. Further taxonomic and ecological work would be necessary to better differentiate between truly local and potentially migratory species within this group. These findings highlight the importance of considering both overwintering strategy and migratory behavior when assessing pest pressure and developing monitoring or management tools. As climate conditions continue to shift, the phenological responses of both local and migratory species may become increasingly asynchronous, complicating prediction models and requiring adaptive strategies in pest management.

We observed temporal shifts in leafhopper community composition, as indicated by diversity and abundance indices, when combining our dataset from 2021 to 2024 (Plante et al. 2024). Of the 69 genera recorded, 44 were consistently observed throughout the study period. The number of unobserved genera progressively declined, from 17 in 2021 and 18 in 2022, to 10 in 2023, and just 5 in 2024. The Hill-Shannon index (which weights genera by frequency) peaked in 2022, whereas the Hill-Simpson index (which emphasizes dominant genera) reached its highest value in 2024. This suggests a decreasing dominance structure in the community over time. To our knowledge, this is the first long-term study of leafhopper diversity in strawberry fields in Québec and worldwide. In contrast, a previous survey in provincial vineyards identified only 11 genera between 2007 and 2008 (Saguez et al. 2014). These initial findings suggest an increase in the number of genera present in strawberry fields. However, these results should be interpreted with caution due to potential differences in host plants, sampling methodologies, temporal scale, and landscape composition, all of which can influence leafhopper community structure (Helbing et al. 2017). Nonetheless, the newly detected genera are of particular interest, especially those associated with known vectors of plant pathogens.

Of particular concern is the first confirmed detection of *Dalbulus maidis* in Canada, a significant pest in South, Central and part of North America, and a known vector of multiple maize pathogens, including *Maize bushy stunt phytoplasma* and corn stunt spiroplasma *Spiroplasma kunkelii* (Nault and Bradfute 1979; Tsai, 2008; Pérez-López et al. 2016; Duffeck et al. 2025). This species has permanent populations in Mexico and further south in the Americas but occasionally irrupts into the United States where it can cause outbreaks of corn stunt spiroplasma (Duffeck et al. 2025). The largest irruption seen to date was in 2024, where it was widely reported across the central and midwestern United States (Faris et al. 2024, Lagos-Kutz et al. 2025). Increased populations in these areas may have contributed to the arrival of the species in Canada.

While strawberries are not a known host for this species, the proximity of corn fields might be the reason why we have detected this pest. The appearance of *D. maidis* highlights the growing role of long-distance dispersal in shaping leafhopper communities in temperate regions and underscores the urgent need for proactive monitoring strategies. The specimens were collected in late September 2024, raising the possibility that their arrival was linked to extreme meteorological events, specifically Hurricane Helene, which moved through Florida and Georgia as a Category 4 storm with sustained winds over 220 Km/h (Meng et al. 2025). We hypothesize that strong wind currents associated with this event may have carried individuals northward into Québec. However, this scenario remains unconfirmed. Alternative explanations, such as the presence of undetected local populations or the establishment of phytoplasma reservoirs, must also be considered. Ongoing monitoring and pathogen screening will be essential to clarify the establishment status of *D. maidis* and its potential epidemiological impact in Canada. Additionally, the detection of *Japananus hyalinus*, a maple specialist, further illustrates the incidental capture of non-crop-associated species. This adventive species has been established in North America since 1897 and in Ontario since 1957 (Hamilton 1983), but its recent detection in Québec highlights the importance of understanding the climatic drivers, dispersal mechanisms, and phenological patterns that influence the movement and establishment of invasive members of the subfamily Auchenorrhyncha in northern agroecosystems (Guglielmino et al. 2002; 2013).

Ultimately, our findings provide a baseline for understanding how climate-sensitive traits, such as overwintering biology, temperature thresholds, and precipitation responses, shape the dynamics of migratory and local leafhoppers. They also emphasize the potential for northward expansions of pest species previously confined to warmer regions, which may introduce new risks to Canadian agriculture. These insights are essential for the development of climate-resilient pest monitoring systems. They lay the groundwork for tailored, species-specific strategies in integrated pest management, including adjusted monitoring schedules, phenology-based forecasting, and targeted interventions. Such precision approaches will be critical to reduce reliance on broad-spectrum synthetic insecticides while maintaining crop productivity and preserving insect biodiversity under a rapidly changing climate.

The need for alternative and sustainable pest control strategies is becoming increasingly urgent. Our field data confirm that, as previously reported (Plante et al. 2024), insecticides currently used by strawberry growers in Québec fail to reduce leafhopper abundance. Despite routine applications, migratory and local leafhopper abundance remained stable before and after intervention across farms, mirroring trends first observed during the 2021–2022 sampling seasons (**Fig. S7, Table S19**). This ongoing inefficacy of chemical control, the lack of products specifically designed to control leafhoppers, combined with climate-driven increases in leafhopper abundance and voltinism, underscores the urgent need to rethink current pest management practices and prioritize the development of sustainable alternatives.

One promising alternative for sustainable pest management lies in increasing the biological control by parasitoid wasps. They have long been recognized for their role in regulating leafhopper populations in North American regions such as Mexico (Moya-Raygoza and Becerra-Chiron, 2014; Torres-Moreno and Moya-Raygoza, 2020) but remain virtually uncharacterized in Canada. Among these, members of the Dryinidae family are particularly noteworthy. Unlike egg parasitoids in the Mymaridae (e.g., *Anagrus* spp.), which oviposit in leafhopper eggs and target species such as *E. fabae* (Triapitsyn et al. 2019) and *D. maidis* (Moya-Raygoza et al. 2012), Dryinidae parasitize the nymphal and adult stages (Olmi and Virla, 2014). This biological difference is crucial, as it makes yellow sticky traps, used during our survey, an effective method for detecting Dryinid parasitism and evaluating its seasonal and environmental determinants influencing it.

Our findings provide the first ecological and genomic characterization of Dryinidae parasitoids, specifically *Gonatopus* spp., within Canadian cropping systems. Parasitism rates averaged around 3%. While the average rate may seem low, an important point to mention is that yellow sticky traps mostly attract adult leafhoppers. As a result, the abundance of parasitized nymphs could not be properly evaluated, likely leading to an underestimation of the actual parasitism rate in the field. Since leafhopper nymphs do not fly and move less than adults, they may be better hosts for Dryinidae parasitoids, and the proportion of parasitized individuals among them could be relatively high. Also, parasitism rates exceed 20% for several weeks, showing a seasonal trend.

Rates increased significantly with temperature and Julian day, peaking in late summer and early fall. The migratory leafhopper *M. quadrilineatus* accounted for more than 80% of parasitized individuals, with peak numbers from July to September. In contrast, *E. fabae*, despite being the second highest in abundance, exhibited minimal parasitism, an observation potentially explained by differences in phenology, behavior, or host suitability (Hoy et al. 1992; Santos et al. 2025). Precipitation, in contrast to temperature, did not have a significant influence on parasitism occurrence. These trends suggest that *Gonatopus* parasitoids in Eastern Canada are primarily constrained by thermal and seasonal factors rather than rainfall, and that warming conditions may, paradoxically, enhance their activity and biocontrol effectiveness in temperate agroecosystems.

This is particularly relevant considering the recent invasion of North America by the Spotted Lanternfly (*Lycorma delicatula*), a polyphagous planthopper from Asia first detected in the U.S. in 2014 (Barringer et al. 2014) and already intercepted multiple times in Canada (CFIA, 2024). The case of *L. delicatula* underscores the urgency of identifying native or introduced natural enemies capable of responding to invasive species. In China, *Dryinus sinicus*, a dryinid wasp with strong host specificity, has been shown to parasitize lanternfly nymphs at rates of up to 30% (Xin et al. 2020; Urban and Leach, 2023; Zhuang et al. 2025). Although *D. sinicus* has not been introduced into North America, our findings demonstrate that native dryinids, such as *Gonatopus*, are already active and seasonally synchronized with pest populations, suggesting promising potential for conservation biocontrol in response to future incursions.

Equally novel are our integrative taxonomic results, which mark a significant advancement in dryinid systematics. We identified eleven *Gonatopus* specimens from Québec, using both classical morphological keys (Olmi, 1984) and comparative material from the Canadian National Collection. Morphological analyses revealed three distinct morphotypes, including one matching *Gonatopus clavipes*, formally recorded in Québec and Canada for the first time. Although *G. clavipes* had been listed as a GBIF occurrence from Manitoba and Nunavut, no peer-reviewed confirmation existed until now. Two other morphotypes displayed traits inconsistent with any described species, suggesting the presence of at least - two potentially undescribed Nearctic taxon.

Our genomic analysis further supported this diversity. Using ancient and degraded DNA workflows, we reconstructed complete or near complete mitogenomes from two parasitoid specimens and one from parasitized M. *quadrilineatus*, while producing *coxI* data for nine of the adults and one larva. Notably, we report the first full mitogenome of a *Gonatopus* species from the New World, the only currently available, although the genome of *G. flavifemur* from China is available (Yang et al. 2021). The mitogenome derived from a *Macrosteles*-associated larva was 16,502 bp in length and featured canonical insect mitochondrial gene content and structure. Phylogenomic analysis based on *coxI* and whole-mitogenome alignments revealed three well-supported clades: one matching *G. clavipes* with >99% identity, one exclusive to *M. quadrilineatus*-associated individuals, which support that this species is actively reproducing in the field as we found here adults, larva and parasitized leafhoppers, and finally one clade containing seven specimens lacking close matches in public databases. These results show congruence between morphological and molecular data, providing an integrative framework for the future of dryinid taxonomy in Canada. If to all this we add that we also found that *Gonatopus* specimens are feeding from a diverse array of leafhoppers, we have evidence that they could be strong candidates to control migratory and local populations.

Unexpectedly, our genomic analyses revealed a third critical trophic component: the entomopathogenic fungus *Beauveria bassiana*, with partial mitogenomes recovered from three Dryinidae specimens and visible mycelial growth in at least two, although hard to say that this mycelium was indeed *B. bassiana*. The fungi *B. bassiana* is a widely used microbial control agent effective against numerous Hemipteran pests, including leafhoppers and planthoppers, where it frequently achieves 60–90% mortality in species such as *Empoasca vitis* and *Nilaparvata lugens*, both in laboratory and field settings (Pu et al. 2005; Erper et al. 2022). These findings open new research avenues for integrated pest management. On one hand, *B. bassiana* could act in conjunction with Dryinidae parasitism to intensify leafhopper control. On the other hand, fungal infection might reduce parasitoid fitness or disrupt host–parasitoid dynamics. Experimental approaches, such as assessing spore viability on parasitized hosts and conducting co-infection assays, are necessary to discern synergistic or antagonistic outcomes.

## Future directions and implications of the study

This study underscores how climate, species biology, ecology, and movement are reshaping pest dynamics in Eastern Canada’s agroecosystems. The increasing abundance of migratory leafhoppers, combined with the first confirmed detection of *D. maidis* in Canada, suggests a future where long-distance dispersal events, likely intensified by extreme weather, will become more frequent. As climate thresholds for population growth between migratory and locals become clearer, such as the sharp increases in *E. fabae* and *M. quadrilineatus* beyond 14 to 16°C, this information can improve phenology-based forecasts to anticipate outbreaks.

Equally important, our study shows the potential for conservative biological control in reducing the population size of leafhoppers. Our findings reveal, for the first time, that native *Gonatopus* parasitoids act in both migratory and local species, and parasitism rates rise in late summer positively with temperature increases. The use of genomic tools confirmed parasitoid diversity, including a first New World mitogenome for *Gonatopus*, and opens new avenues for taxonomic resolution and tracking biological control agents. Together, these insights provide a foundation for building a more ecologically sound pest management systems that respond to changing climates while minimizing reliance on synthetic insecticides.

## DATA AVAILABILITY

All genomic data generated in this study are publicly available in the NCBI database under BioProject [PRJNA1294321]. This submission includes 15 BioSamples, comprising 11 derived from ancient DNA sequencing workflows, 3 from Illumina sequencing, and 1 *cox1* sequence from larval material. A total of 14 Sequence Read Archive (SRA) datasets are available, along with 3 complete mitochondrial genomes and 1 *cox1* gene sequence. The full dataset can be accessed at: [https://www.ncbi.nlm.nih.gov/bioproject/PRJNA1294321]. One specimen of *Dalbulus maidis* was deposited into the Canadian National Collection of Insects, Arachnids, and Nematodes under voucher number CNC2151743.

## CODE AVAILABILITY

Codes are available from GitHub: https://github.com/Edelab

## AUTHOR CONTRIBUTIONS

J.J., A.A.S., and N.P. contributed to data collection, insect sorting and identification. J.J., and A.A.S. led the formal analysis, visualization, and interpretation of ecological data. J.J., J.M., and F.S. performed DNA extractions and contributed to bioinformatic analyses. J.M., and F.S. conducted molecular identification and genome assemblies. N.P. coordinated fieldwork, supported leafhopper identification and supported methodology development. J.L.F.T., J.K., J.M.D-S. and J.-F.L. assisted in the taxonomic identification of parasitoid specimens and provided access to reference collections. F.M. contributed to climate modeling and statistical support. V.F. provided guidance on experimental design and supported interpretation within an agroecological context, and secured funding. E.P.-L. conceptualized the study, secured funding, supervised the project, and coordinated manuscript preparation. J.J., A.A.S. and E.P.-L. wrote the original draft, with all authors contributing to review and editing.

## ACKNOWLEDGMENTS

This work was supported by RQRAD, MAPAQ, and FRQNT through the *Programme de recherche en partenariat*—*Agriculture durable*—*Volet II*—*2e concours* (application no. 337847), as well as by NSERC through the Alliance-SARI Program (Grant ALLRP 588519-23). We thank Arthur Thompson de la Chenelière, Myriam Moreault, Marie-Fleur Dallaire, Mathilde Bélanger, and Gabrielle Le Gal for their contributions to insect sorting and identification. We are also grateful to Valeria Mejia Ruiz, supported by the NSERC-USRA program, to Chenyu Huang, supported by the MITACS Globalink program, and to Jordanne Jacques supported through travelling scholarships by QCSB and RQRAD. Finally, we extend our sincere thanks to our long-time collaborators, the strawberry growers and the dedicated personnel at their farms for their continued support and participation.

## ETHICS

This work complied with the relevant legal requirements of the Université Laval. The leafhopper or parasitoid species studied here are not endangered species.

## ADDITIONAL INFORMATION

### Supplementary Tables

**Table S1**. Detailed information on the strawberry fields included in this study.

**Table S2**. Model selection and diagnostics for GLMMs assessing seasonal variation in leafhopper abundance.

**Table S3**. Model selection and diagnostics for GLMMs evaluating the effect of climatic variables on leafhopper abundance.

**Table S4**. Detailed information on the insecticides used by strawberry growers during both growing seasons included in the study, and those treatments were selected for further statistical analyses.

**Table S5**. Model selection and diagnostics for GLMMs evaluating monthly parasitism rates (beta-binomial models with spline transformations) and parasitism occurrence, including effects of time of the season, precipitation, and temperature.

**Table S6**. Detailed information of adult Dryinidae specimens captured in this study.

**Table S7**. Detailed information on *Macrosteles quadrilineatus* individuals parasitized by *Gonatopus* spp. and analyzed using Illumina sequencing.

**Table S8**. Detailed information of the *Gonatopus* spp. and the *coxI* GenBank accession numbers used in the phylogenetic analysis.

**Table S9**. Comprehensive list of leafhopper genera recorded by region and year in strawberry fields across Québec, Canada.

**Table S10**. Detailed information of *Dalbulus maidis* specimens captured in this study.

**Table S11**. Detailed summary of diversity index calculations and associated statistical analyses.

**Table S12**. Annual parasitism rates recorded for each leafhopper species throughout the study period.

**Table S13**. Comprehensive feature table of the mitochondrial genome assembled from parasitized *Macrosteles quadrilineatus* specimens, including all protein-coding genes and tRNA annotations with their respective positions, strands, and start codons.

**Table S14**. Summary of sequencing results from ancient and degraded DNA workflows obtained from adult *Gonatopus* parasitoids and one larval specimen.

**Table S15**. Comprehensive feature table of the mitochondrial genome assembled from specimen 10, including all protein-coding genes and tRNA annotations with their respective positions, strands, and start codons.

**Table S16**. Comprehensive feature table of the mitochondrial genome assembled from specimen 5, including all protein-coding genes and tRNA annotations with their respective positions, strands, and start codons.

**Table S17**. Gene length and percentage of missing data for the mitochondrial genes (*atp6, atp8, cob, cox1, cox2, cox3, nad1, nad2, nad3, nad4, nad4l, nad5*, and *nad6*) used in the phylogenetic reconstruction shown in **Figure S6**.

**Table S18**. Summary of host DNA barcoding results based on cox1 sequences recovered from adult *Gonatopus specimens*.

**Table S19**. Details of the specimens from which Beauveria bassiana sequences were recovered.

**Figure S1. Top five most abundant leafhopper genera from 2021 to 2024.** *Macrosteles* and *Empoasca* consistently ranked as the top two genera, followed by *Graminella* in third place across all years. *Hebata* appeared among the top five in three of the four years, while *Paraphlepsius* was among the top five in the last two years.

**Figure S2. Effectiveness of insecticides on leafhopper population control.** Impact of insecticides commonly used by strawberry growers on leafhopper abundance. The nine insecticides applied more than three times between 2023 and 2024, and effects were evaluated across three leafhopper groups: the migratory species *E. fabae* (green) and *M. quadrilineatus* (orange), as well as local species (blue). Only acetamiprid showed a significant reduction in the abundance of *E. fabae* and local species. The remaining insecticides did not lead to significant population decreases. Detailed data of the products used is presented in **Table S4**.

**Figure S3.** Migratory and local leafhoppers with visible ectoparasitoid larval sacs (indicated by white arrows). (**a**) *Macrosteles quadrilineatus*, the most frequently parasitized migratory species. (**b**) *Graminella nigrifrons*, the most frequently parasitized local species detected in this study.

**Figure S4.** Morphological characterization of *Gonatopus* specimen 11. (**a**) Close-up view of the chelate protarsi, showing what appears to be a single row of lamellae. (**b**) Frontal view and (**c**) dorsal view of specimen 11. This individual was retrieved from a yellow sticky trap, which explains its degraded condition. Molecular analyses suggest that this specimen corresponds to the species parasitizing *Macrosteles quadrilineatus* in Eastern Canada.

**Figure S5.** Morphological characterization of *Gonatopus* specimen 10. (**a–b**) Close-up views of the chelate protarsi, showing a single row of 21 lamellae on the distal half of protarsomere V. (**c**) Dorsal view of specimen 10. This specimen was identified based on morphology and molecular evidence as *Gonatopus clavipens*. The white arrow points to mycelium possibly belonging to Beauveria bassiana.

**Figure S6.** Maximum likelihood phylogenetic tree using the mitochondrial genes *atp6, atp8, cob, cox1, cox2, cox3, nad1, nad2, nad3, nad4, nad4l, nad5*, and *nad6* concatenated. The scale bar represents 1 substitution per 10 nucleotide positions. Further details about the genes are provided in **Table S17**.

**Figure S7.** Mitochondrial genome reconstruction of Beauveria bassiana from specimen *7*. (**a**) Newly assembled *B. bassiana* mitogenome, with the red segment indicating a region missing compared to (**b**) the reference genome of this entomopathogenic fungus. (**c**) Dorsal view of specimen 7. (**d**) Close-up of the chelate protarsi showing a lamellar arrangement like that of the representative specimen shown in **Figure 6**.

