## Supplementary material for "Uncovering diversity and climatic drivers of leafhopper-parasitoid dynamics in Canada": Figure S1 to S7

**Supplementary Figures**

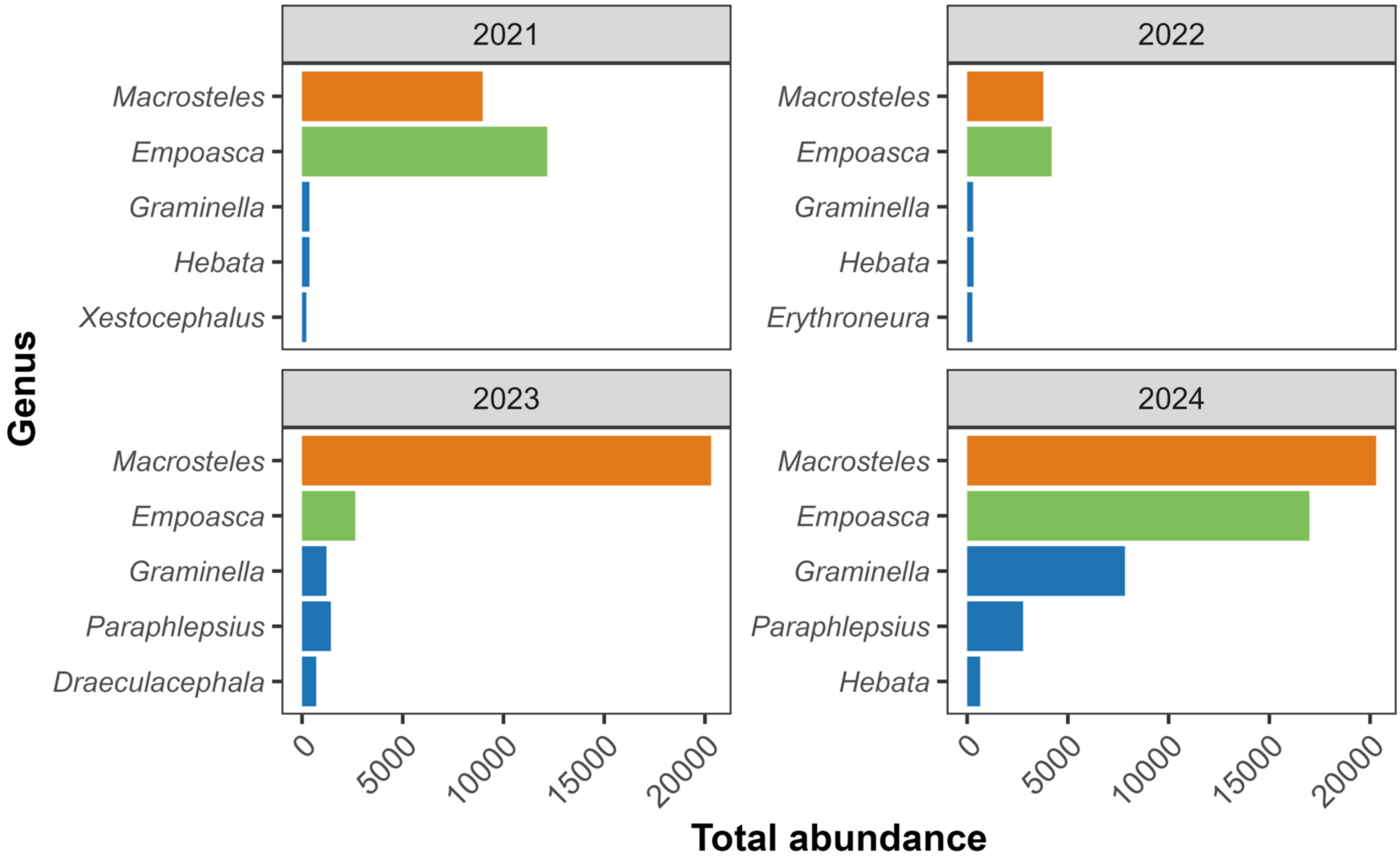


**Fig. S1. Top five most abundant leafhopper genera from 2021 to 2024.** *Macrosteles* and *Empoasca* consistently ranked as the top two genera, followed by *Graminella* in third place across all years. *Hebata* appeared among the top five in three of the four years, while *Paraphlepsius* was among the top five in the last two years.

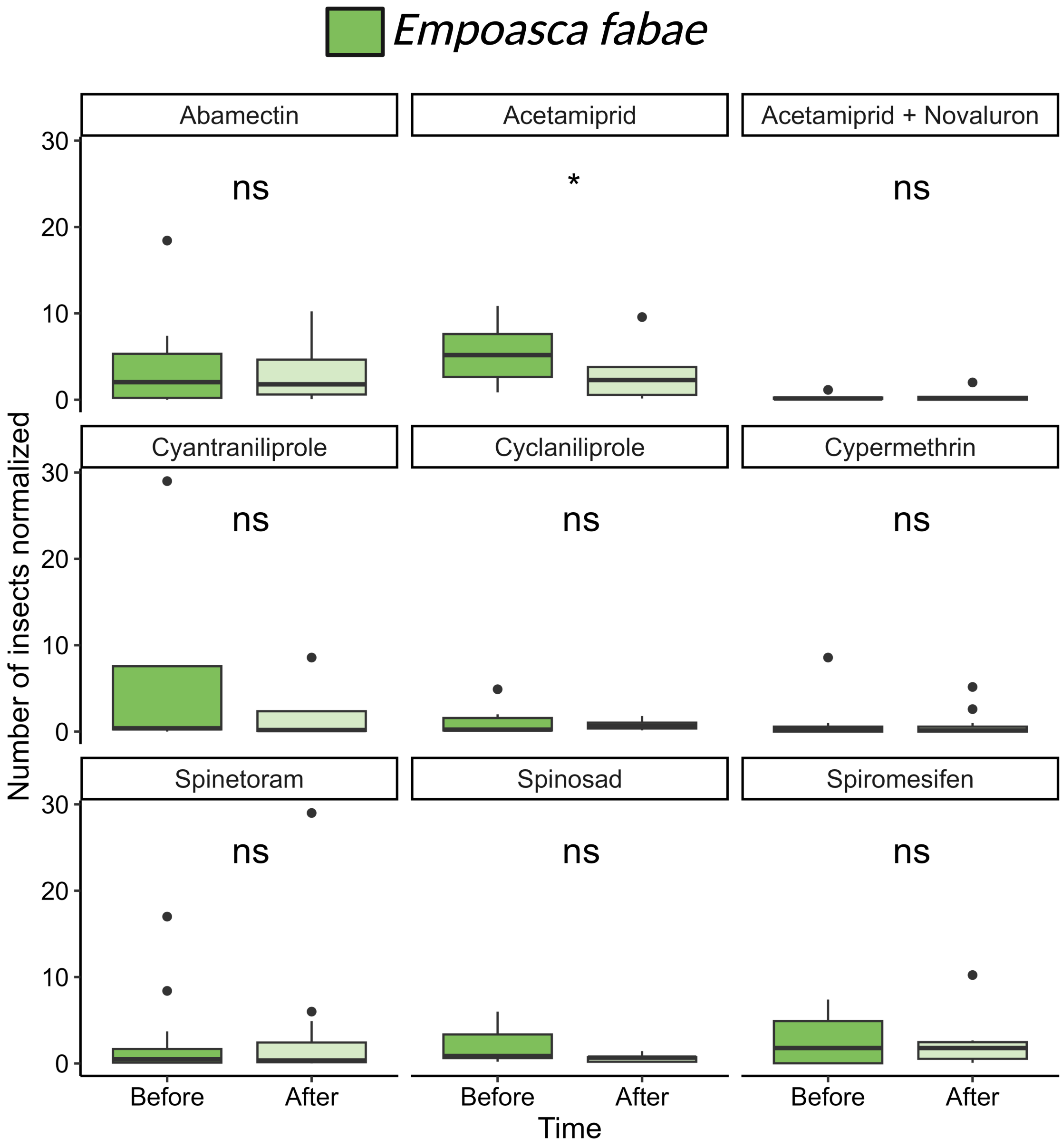

Empoasca fabae
Abamectin
Acetamiprid
Acetamiprid + Novaluron
Cyantraniliprole
Cyclaniliprole
Cypermethrin
Spinetoram
Spinosad
Spiromesifen
ns
*
Number of insects normalized
0
10
20
30
Before
After
Time

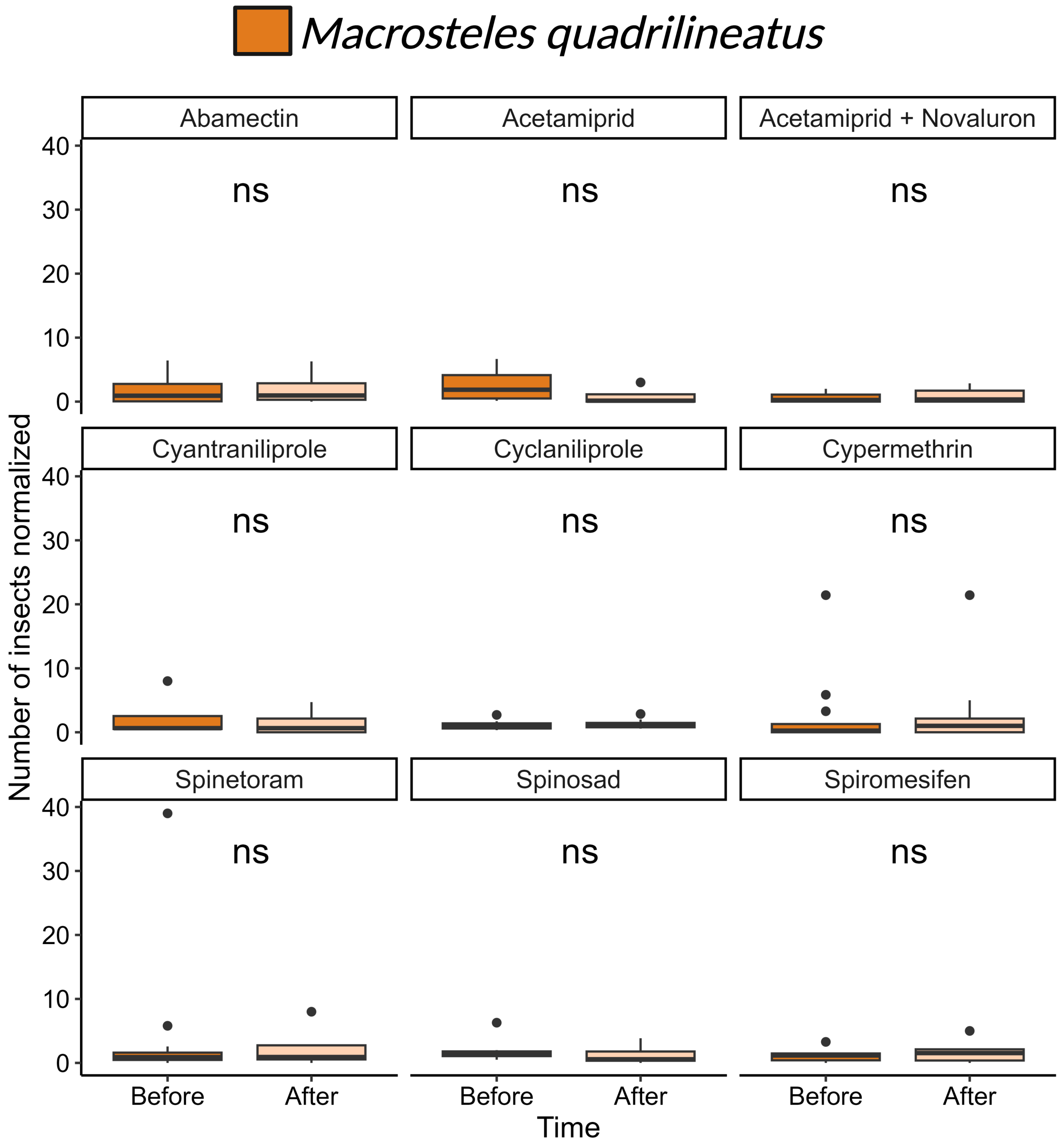
Macrosteles quadrilineatus
Abamectin
Acetamiprid
Acetamiprid + Novaluron
Cyantraniliprole
Cyclaniliprole
Cypermethrin
Spinetoram
Spinosad
Spiromesifen
ns
Number of insects normalized
0
10
20
30
40
Before
After
Time

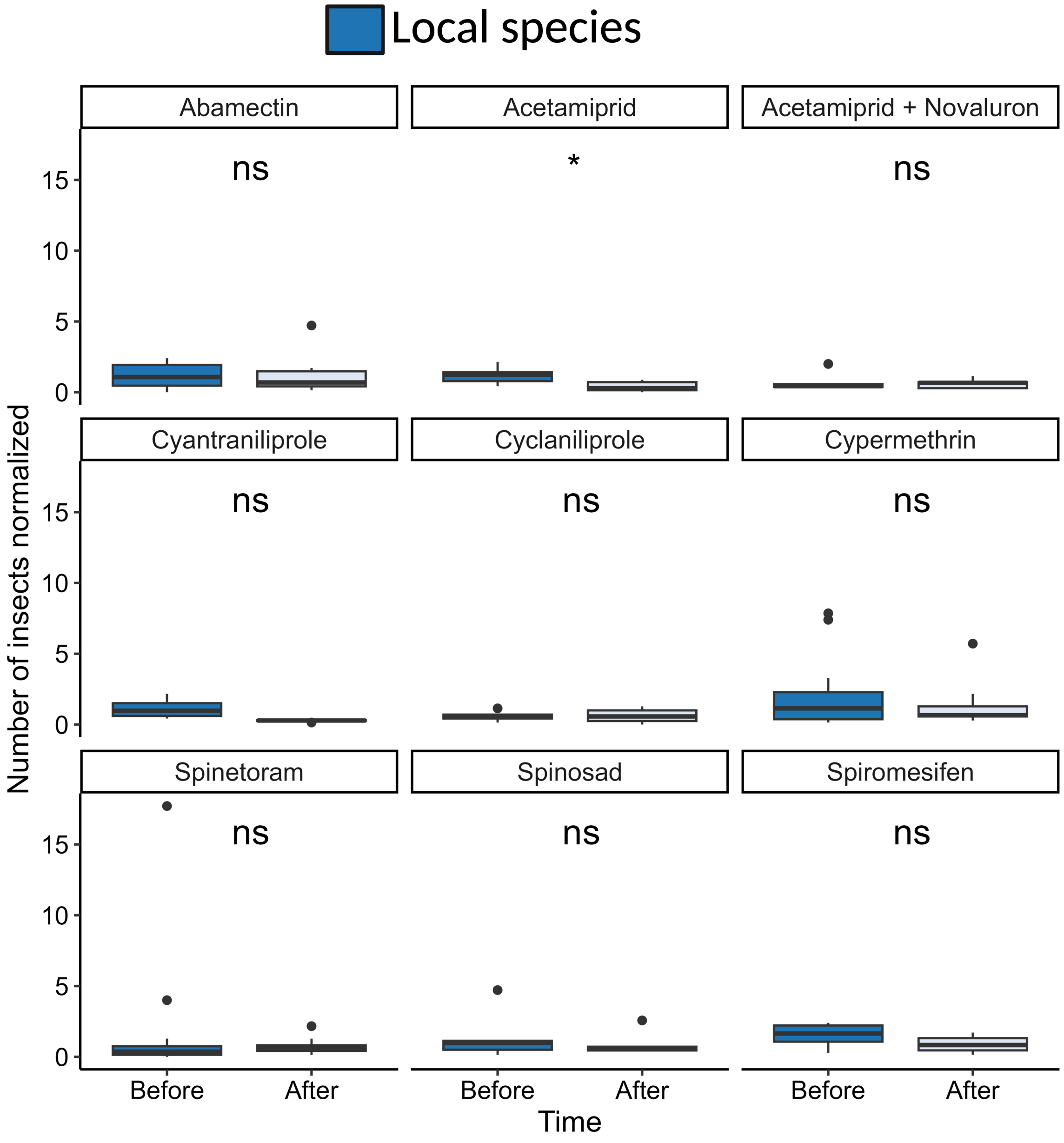


**Fig. S2. Effectiveness of insecticides on leafhopper population control.** Impact of insecticides commonly used by strawberry growers on leafhopper abundance. The nine insecticides applied more than three times between 2023 and 2024, and effects were evaluated across three leafhopper groups: the migratory species *E. fabae* (green) and *M. quadrilineatus* (orange), as well as local species (blue). Only acetamiprid showed a significant reduction in the abundance of *E. fabae* and local species. The remaining insecticides did not lead to significant population decreases. Detailed data of the products used is presented in **Table S4**.

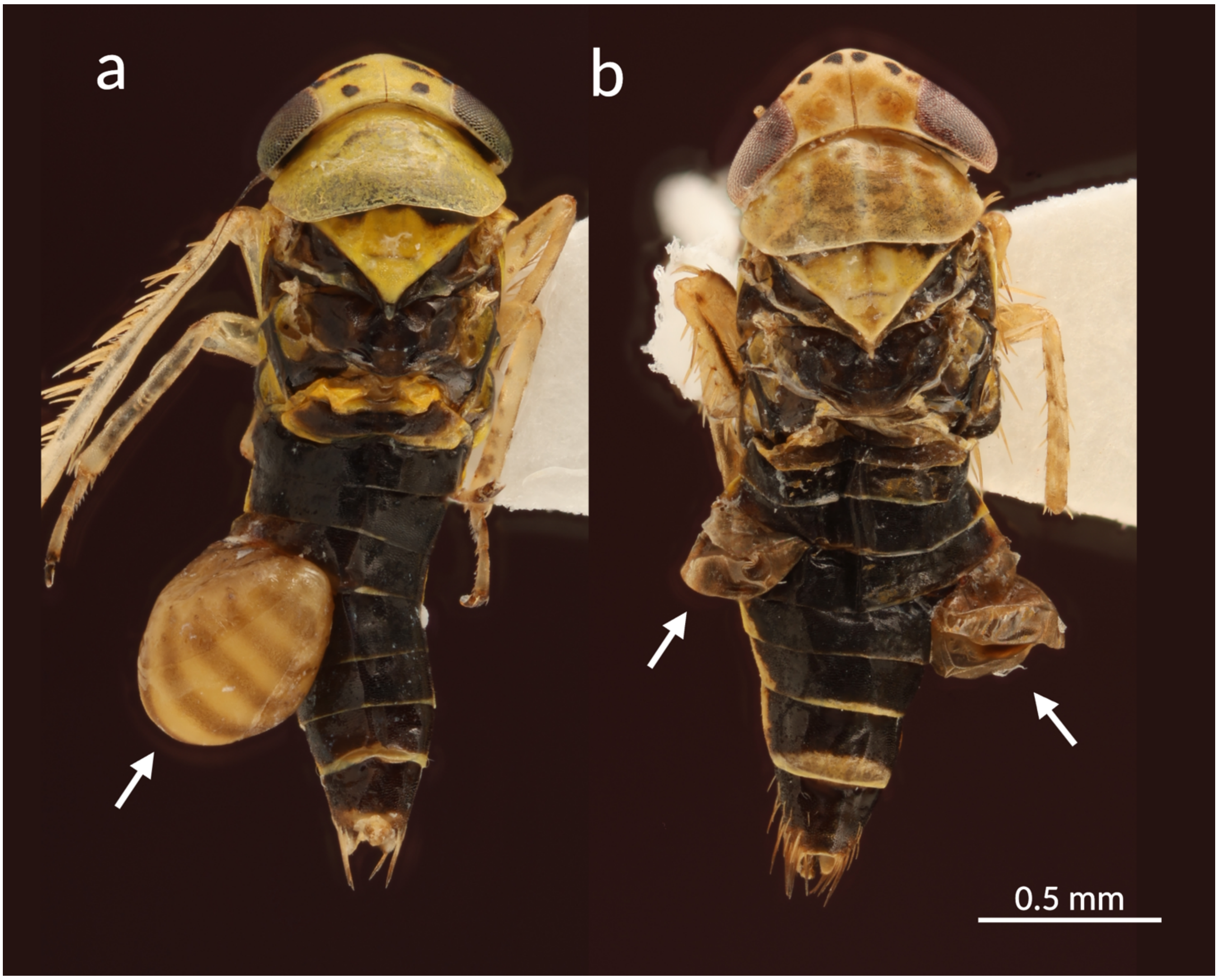


**Fig. S3.** Migratory and local leafhoppers with visible ectoparasitoid larval sacs (indicated by white arrows). (**a**) *Macrosteles quadrilineatus*, the most frequently parasitized migratory species. (**b**) *Graminella nigrifrons*, the most frequently parasitized local species detected in this study.

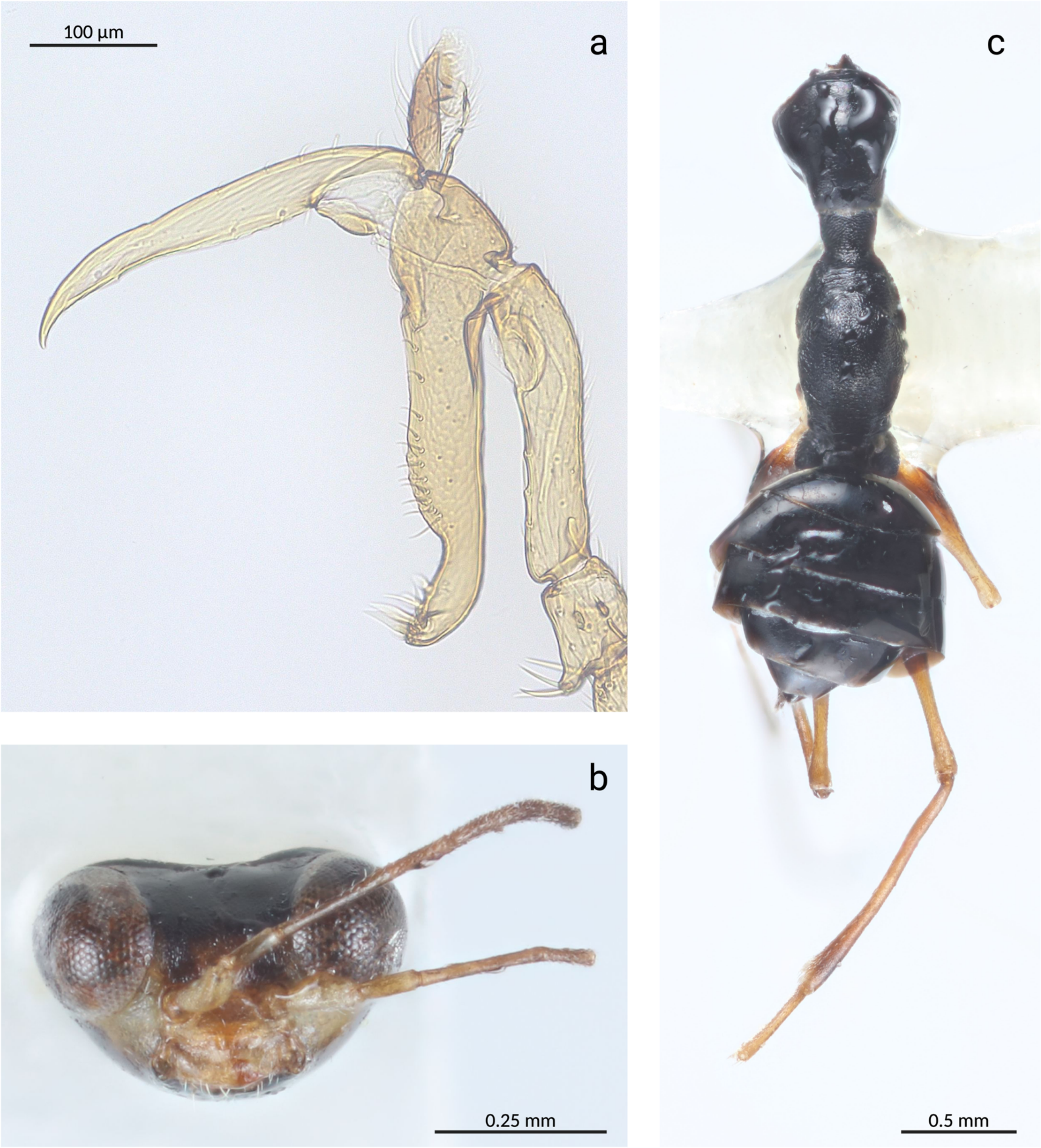


**Fig. S4.** Morphological characterization of *Gonatopus* specimen 11. (**a**) Close-up view of the chelate protarsi, showing what appears to be a single row of lamellae. (**b**) Frontal view and (**c**) dorsal view of specimen 11. This individual was retrieved from a yellow sticky trap, which explains its degraded condition. Molecular analyses suggest that this specimen corresponds to the species parasitizing *Macrosteles quadrilineatus* in Eastern Canada.

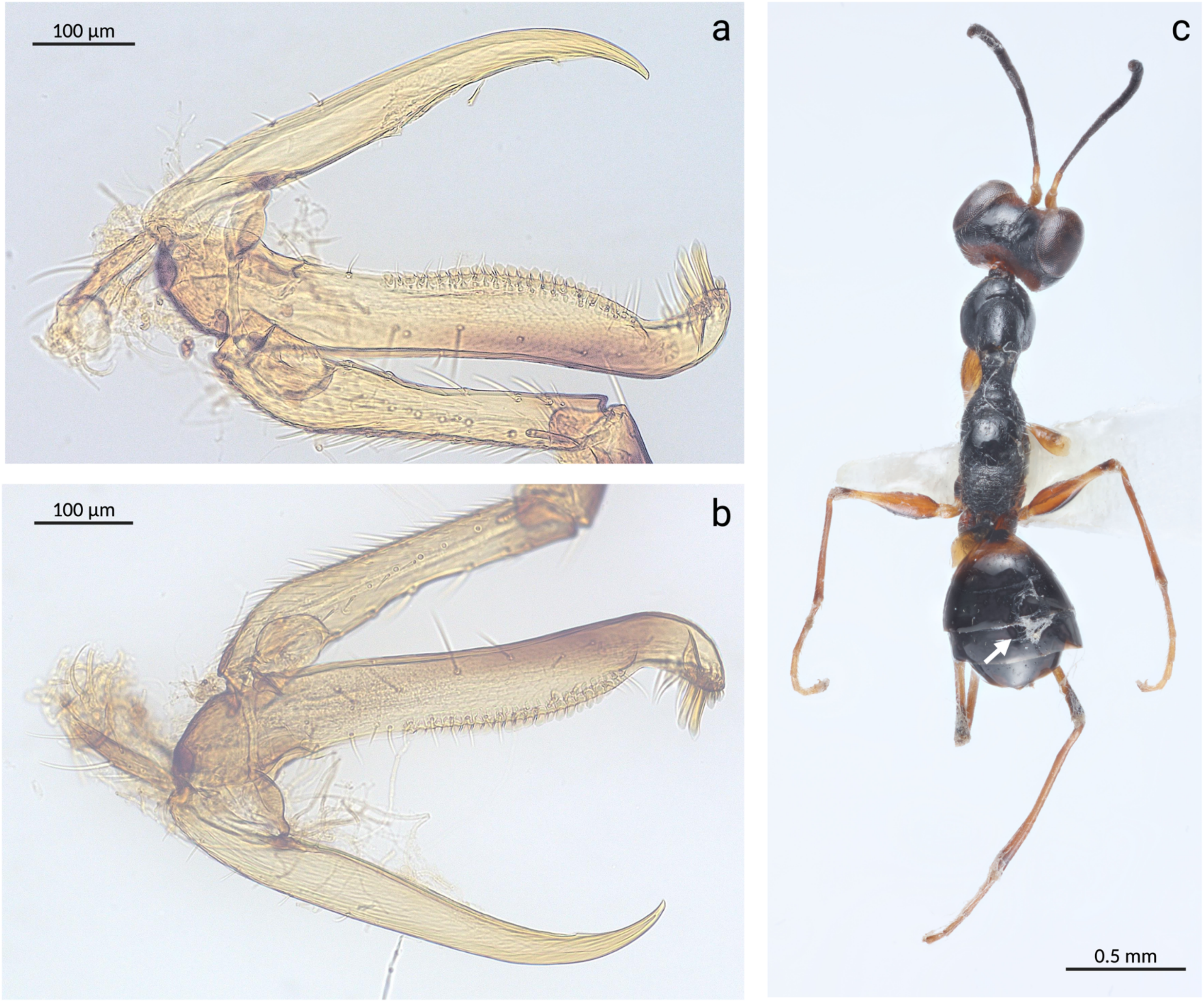


**Fig. S5.** Morphological characterization of *Gonatopus* specimen 10. (**a–b**) Close-up views of the chelate protarsi, showing a single row of 21 lamellae on the distal half of protarsomere V. (**c**) Dorsal view of specimen 10. This specimen was identified based on morphology and molecular evidence as *Gonatopus clavipens*. The white arrow points to mycelium possibly belonging to *Beauveria bassiana*.

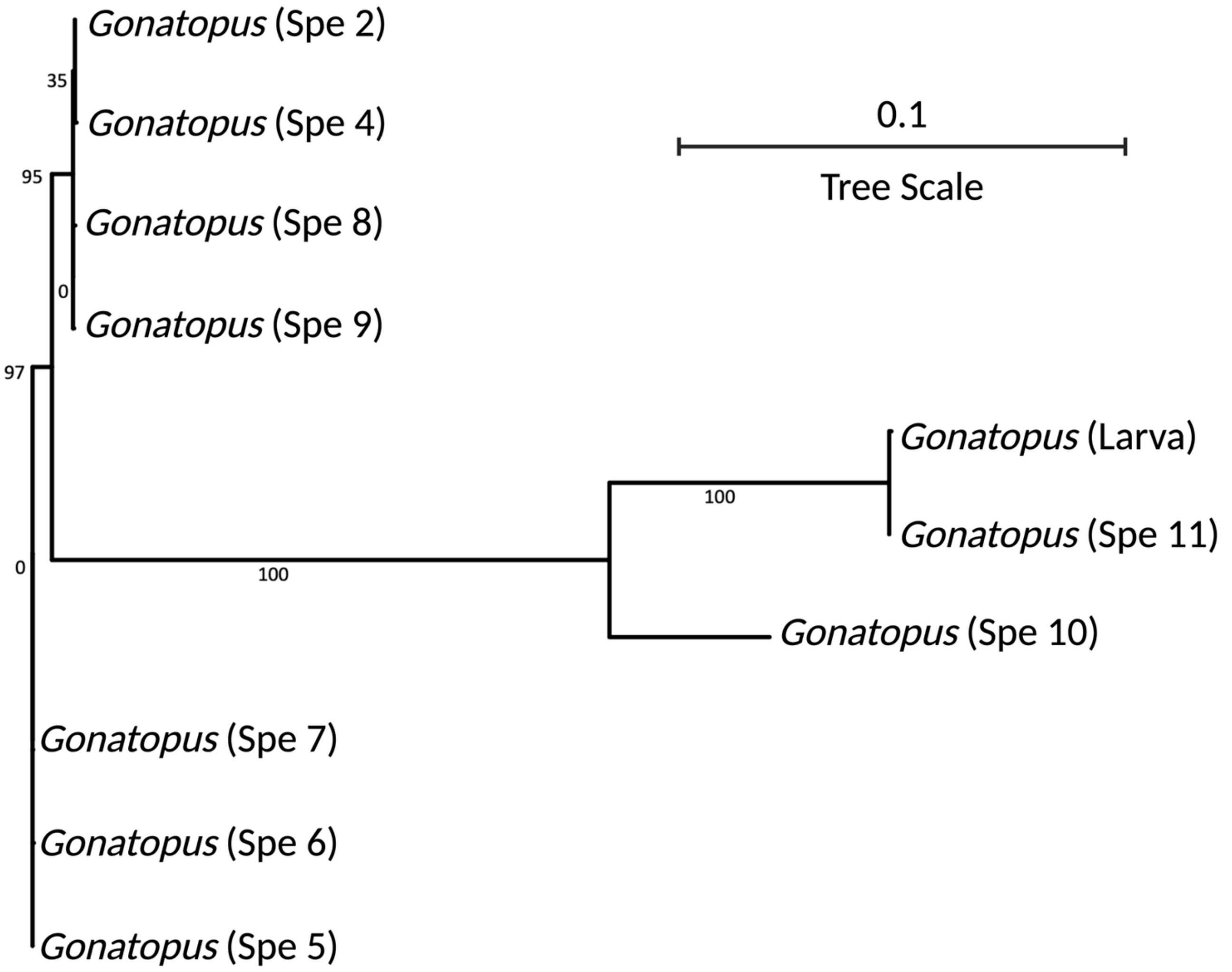


**Fig. S6.** Maximum likelihood phylogenetic tree using the mitochondrial genes *atp6, atp8, cob, cox1, cox2, cox3, nad1, nad2, nad3, nad4, nad4l, nad5,* and *nad6* concatenated. The scale bar represents 1 substitution per 10 nucleotide positions. Further details about the genes are provided in **Table S17**.

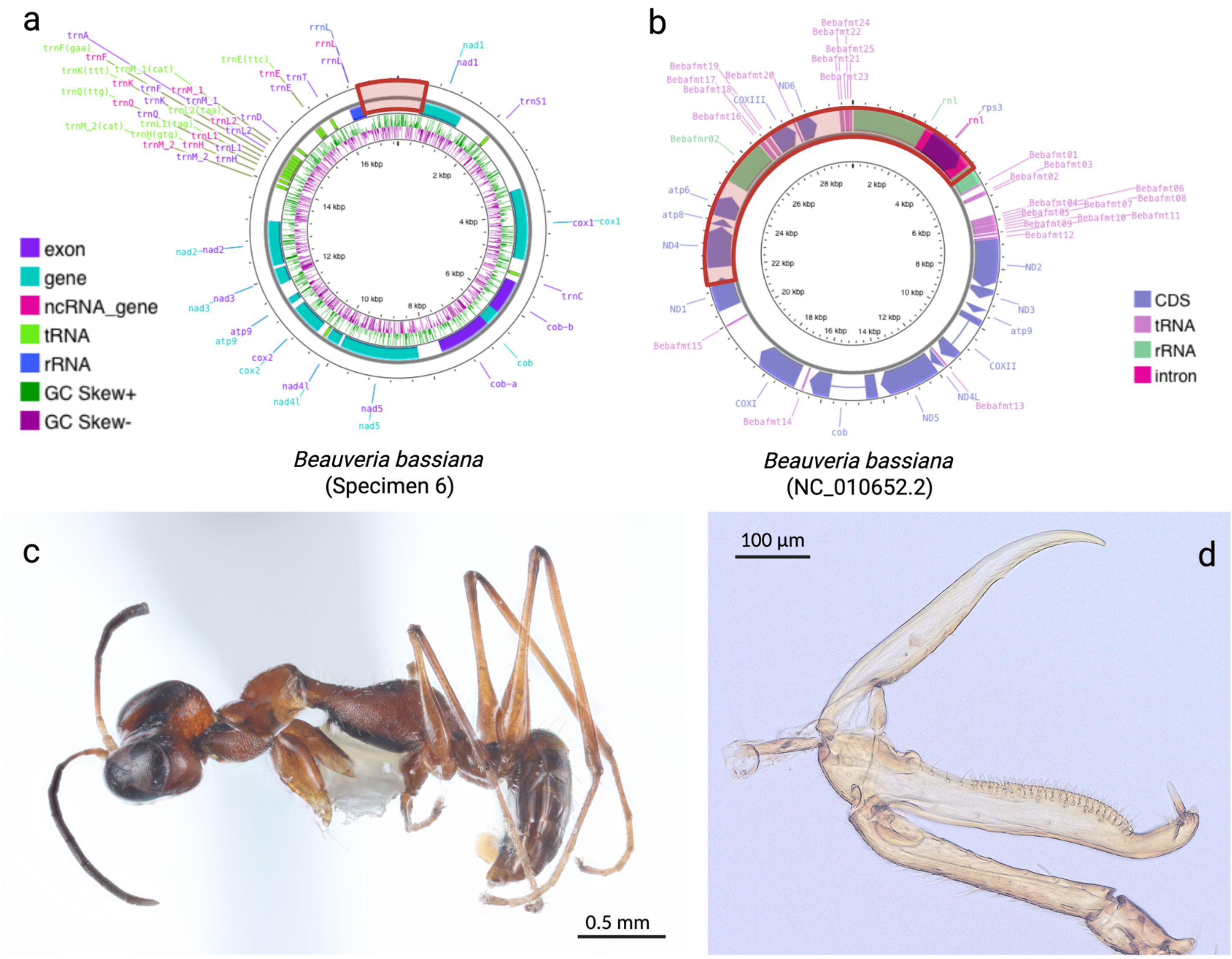


**Fig. S7.** *Mitochondrial genome reconstruction of Beauveria bassiana from specimen 7.* (**a**) Newly assembled *B. bassiana* mitogenome, with the red segment indicating a region missing compared to (**b**) the reference genome of this entomopathogenic fungus. (**c**) Dorsal view of specimen 7. (**d**) Close-up of the chelate protarsi showing a lamellar arrangement like that of the representative specimen shown in **Fig. 6**.
